# Identifying conservation priorities in a defaunated tropical biodiversity hotspot

**DOI:** 10.1101/790766

**Authors:** Andrew Tilker, Jesse F. Abrams, An Nguyen, Lisa Hörig, Jan Axtner, Julie Louvrier, Benjamin M. Rawson, Hoa Anh Nguyen Quang, Francois Guegan, Thanh Van Nguyen, Minh Le, Rahel Sollmann, Andreas Wilting

## Abstract

**Aim:** Unsustainable hunting is leading to widespread defaunation across the tropics. To mitigate against this threat with limited conservation resources, stakeholders must make decisions on where to focus anti-poaching activities. Identifying priority areas in a robust way allows decision-makers to target areas of conservation importance, therefore maximizing the impact of conservation interventions.

**Location:** Annamite mountains, Vietnam and Laos.

**Methods:** We conducted systematic landscape-scale surveys across five study sites (four protected areas, one unprotected area) using camera-trapping and leech-derived environmental DNA. We analyzed detections within a Bayesian multi-species occupancy framework to evaluate species responses to environmental and anthropogenic influences. Species responses were then used to predict occurrence to unsampled regions. We used predicted species richness maps and occurrence of endemic species to identify areas of conservation importance for targeted conservation interventions.

**Results:** Analyses showed that habitat-based covariates were uninformative. Our final model therefore incorporated three anthropogenic covariates as well as elevation, which reflects both ecological and anthropogenic factors. Conservation-priority species tended to found in areas that are more remote now or have been less accessible in the past, and at higher elevations. Predicted species richness was low and broadly similar across the sites, but slightly higher in the more remote site. Occupancy of the three endemic species showed a similar trend.

**Main conclusion:** Identifying spatial patterns of biodiversity in heavily-defaunated landscapes may require novel methodological and analytical approaches. Our results indicate to build robust prediction maps it is beneficial to sample over large spatial scales, use multiple detection methods to increase detections for rare species, include anthropogenic covariates that capture different aspects of hunting pressure, and analyze data within a Bayesian multi-species framework. Our models further suggest that more remote areas should be prioritized for anti-poaching efforts to prevent the loss of rare and endemic species.

## Introduction

Tropical biodiversity is declining at an alarming rate as a result of intense anthropogenic pressures (Bradshaw et al, 2009). Although habitat loss and degradation are major drivers of these declines (Rosa et al, 2016), unsustainable hunting is increasingly emerging as the primary threat to wildlife in tropical biodiversity hotspots (Benítez-López et al, 2017). Large and medium-sized mammals tend to be particularly vulnerable to hunting because they often occur at lower average population densities, have lower intrinsic rates of increase, and longer generation times (Bodmer et al, 1997; Cardillo et al, 2005; Davidson et al, 2009). Indeed the “empty forest syndrome” that Redford (1992) warned about almost three decades ago is now a commonplace phenomenon and, given the ever-increasing demand for wildlife products in the world’s tropical regions (Rosen & Smith, 2010; Ripple et al, 2016), this trend is unlikely to slow in the coming years. Without urgent and effective measures to address overexploitation, tropical wildlife populations will continue to decline, and species extinctions will follow. Confronting the pantropical defaunation crisis has become one of the most important challenges facing conservation today (Bradshaw et al, 2009).

Defaunation has been particularly severe in Southeast Asia, where high human densities, a thriving illegal wildlife trade, weak protected area governance, and rapid infrastructure development have synergistically contributed to unsustainable, industrial-scale hunting (Duckworth et al, 2012; Wilcove et al, 2013; Harrison et al, 2016). Within Southeast Asia, the Annamites ecoregion on the border of Vietnam and Laos has undergone severe defaunation as a result of widespread illegal hunting (Harrison et al, 2016; Timmins et al, 2016). Poaching in the Annamites is primarily accomplished by the setting of indiscriminate wire snares (Gray et al, 2018). Numerous mammals are regionally extinct (Walston et al, 2010; Brook et al, 2014), and even once common species now survive at low densities (Duckworth et al, 2016). High levels of unsustainable hunting pressure are particularly worrisome from a conservation perspective, because the region is home to several endemic mammal species. Mammals restricted to this ecoregion include the saola *Pseudoryx nghetinhensis*, large-antlered muntjac *Muntiacus vuquangensis*, Annamites dark muntjac species complex *Muntiacus rooseveltorum / truongsonensis* Owston’s civet *Chrotogale owstoni*, and Annamite striped rabbit *Nesolagus timminsi* (Tordoff et al, 2003; Hurley et al, 2005; Long et al, 2005). Taken together, the high poaching pressure and unique biodiversity in the Annamites make it one of the highest priority tropical regions in the world for the prevention of imminent hunting-driven extinctions.

To maximize the effectiveness of conservation interventions to prevent unsustainable hunting in tropical biodiversity hotspots, it is imperative to make optimal use of limited conservation resources. In the Annamites, the magnitude of the snaring crisis (Gray et al, 2018), coupled with nascent protected area enforcement capacities and lack of sufficient resources, has overwhelmed efforts to adequately reduce this threat at the landscape level. Given these limitations, targeting snare removal efforts to specific areas within a landscape may be critical to reduce snaring to levels that would allow population recovery. To implement this approach, it is first necessary to identify priority areas. In the Annamites, areas that harbor threatened and endemic species are top priorities for targeted *in situ* protection measures. These species often occur at low densities, and are therefore particularly susceptible to local extirpation. To identify priority areas, it is important to apply appropriate analytical techniques. Species distribution modeling provides an ideal framework for mapping spatial patterns of biodiversity, and thus identifying conservation-priority areas (Rodríguez et al, 2007; Guisan et al, 2013).

There are, however, two fundamental challenges to the modeling of species distributions in tropical rainforest environments. First, tropical mammal species are often difficult to detect because they are rare, elusive, and occur at low densities. Second, even when these species can be detected, it may be difficult to obtain enough data to construct robust species distribution models (Cayuela et al, 2009), particularly in defaunated areas, where mammal populations are depleted.

Advances in noninvasive survey methods and statistical modeling techniques provide ways to address these challenges. Two noninvasive methods have revolutionized surveys for tropical mammals: camera-traps (Tobler et al, 2008) and high-throughput sequencing of environmental DNA (eDNA) (Bohmann et al, 2013). Camera-traps are a well-established method, and have been used to gather data on even the rarest of tropical mammal species (Whitfield, 1998; Raloff, 1999; Ganas & Lindsell, 2010). The use of eDNA is relatively new but shows considerable promise. Invertebrate-derived DNA (iDNA) approaches using terrestrial hematophagous leeches, in particular, have proven adept at detecting tropical mammals (Schnell et al, 2012; Schnell et al, 2018; Weiskopf et al, 2018). Recently, Abrams et al (2019) showed that combining camera-trapping and iDNA leech data has the potential to improve detection probabilities for tropical mammal species beyond what would be provided by each method independently. The joint camera-trap and iDNA approach thus opens new possibilities for obtaining detections of elusive tropical rainforest mammals, which in turn can be used to build robust species distribution models.

Even with improved detection methods and combined datasets, however, it may not be possible to obtain sufficient records for rare species. This shortfall represents a major issue because the rarest species are often the species of highest conservation concern. Multi-species occupancy models offer an analytical framework to address this challenge, as species with few detections borrow information from more abundant species, which allows parameter estimation for rare species (Tobler et al, 2015; Drouilly et al, 2018; Li et al, 2018). Because species-specific responses to covariates can be projected to unsampled areas, this approach can be used to generate maps of species potential occurrence (MacKenzie et al, 2017; Sollmann et al. 2017).

Here, we collected a landscape-scale systematic camera-trapping and iDNA dataset across a protected area complex in the central Annamites landscape to identify priority areas for targeted conservation interventions. We used a multi-species occupancy framework and environmental and anthropogenic covariates to estimate species occurrence and predict species richness across the surveyed landscape. Our prediction maps provide insight into where to focus conservation efforts among individual study sites at the landscape scale, and more specifically can inform deployment of snare-removal teams within protected areas. We discuss our results within the context of informing targeted conservation interventions to prevent further defaunation, and species extinctions, within tropical biodiversity hotspots.

## Methods

### Study area

We conducted landscape-scale surveys in a large contiguous forest in the central Annamites landscape of Vietnam and Laos. The study area spans both countries and is divided into five administrative units. In Vietnam, we surveyed three sites: Bach Ma National Park (NP), the Hue Saola Nature Reserve (NR), and the Quang Nam Saola NR. In Laos, we surveyed the eastern section of Xe Sap National Protected Area (NPA) and an adjacent ungazetted forest near the village of Ban Palé (Fig. 1). Together these areas comprise approximately 900 km^2^ of mountainous terrain with elevations ranging between 100 – 2000 m asl. The dominant habitat type is wet evergreen tropical rainforest. Although the wider central Annamites region has experienced extensive past disturbance from defoliation and logging, habitat loss, degradation, and fragmentation within the past 20 years has been minimal within our study sites (Meyfroidt et al, 2008; Matusch, 2014). At the landscape scale, forest structure and habitat type are consistent across the study sites, characterized by mature secondary forest with a multi-tiered closed canopy (Fig. S1). The Vietnam sites are surrounded by a densely-populated matrix consisting of human settlements, agricultural fields, and timber plantations. Human population density in the Lao sites is low and, aside from small-scale shifting cultivation, the landscape surrounding the survey areas has not been heavily modified. However, Vietnamese incursion into these areas for poaching and illegal gold mining is widespread (Tilker, 2014), and has been facilitated by the recent construction of a road connecting Vietnam and Laos that bisects the Palé area.

**Figure 1:**
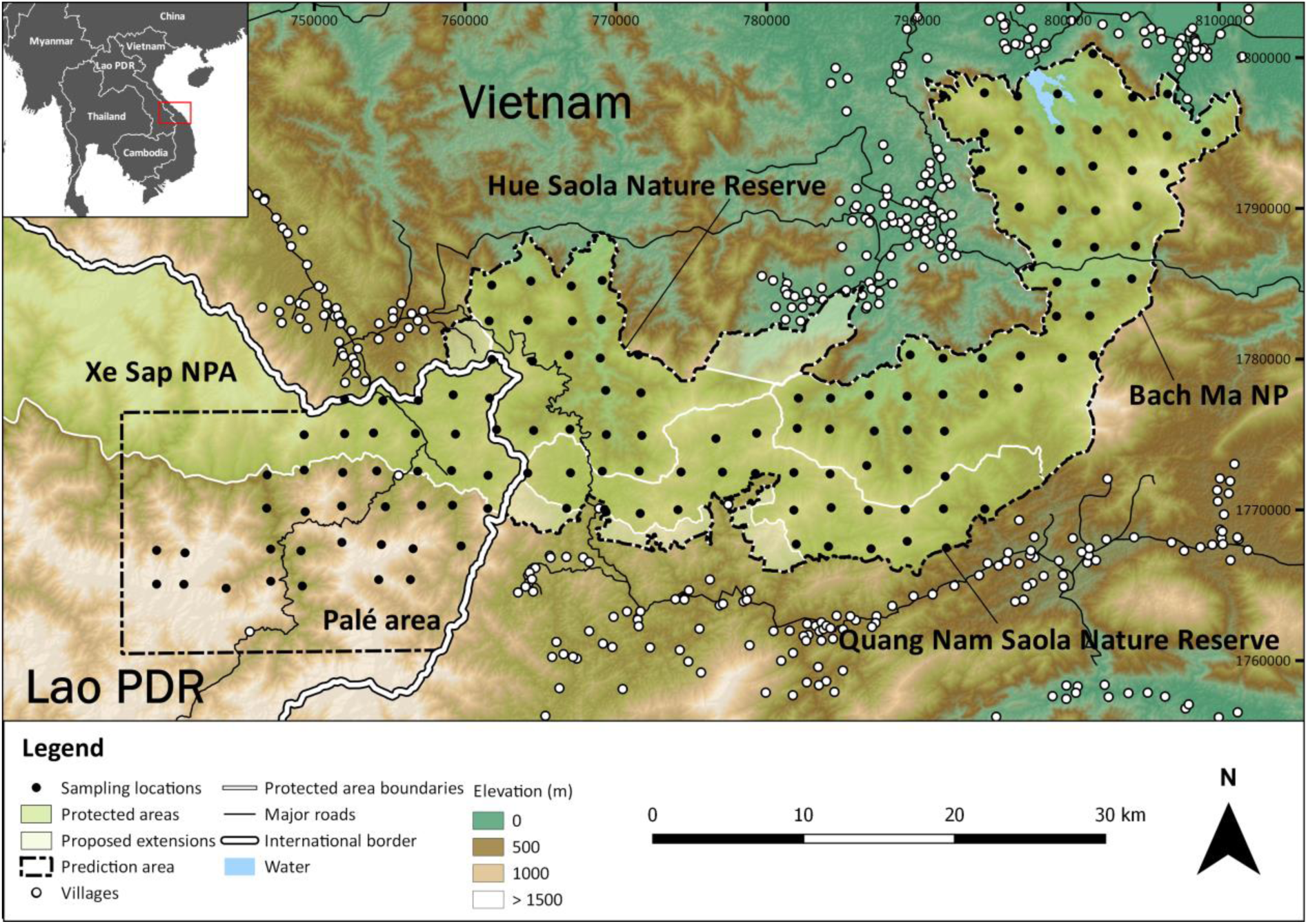
Map of camera-traps and leech collection stations across five areas in the central Annamites landscape.

Poaching pressure is high across the landscape (Wilkinson 2016; WWF, 2017). Measures to mitigate illegal hunting differ in intensity and effectiveness among the five sites. Patrolling in Bach Ma NP is not intensive and has received less technical and financial support than the adjacent sites. The Hue and Quang Nam Saola NRs have benefited from WWF investment in enforcement since 2011 under the Carbon and Biodiversity (CarBi) project, maintaining active Forest Guard patrol teams to strengthen enforcement capacities in the field and provision of capacity development in patrol strategy, data collection and adaptive management for park staff. The Forest Guard teams are comprised of local community members and their primary role is to remove wire snares and destroy poacher camps (Wilkinson, 2017). Between 2011 and 2017, the patrols removed > 110,000 snares from the Hue and Quang Nam Saola NRs (WWF, 2017). The eastern section of Xe Sap NPA has also benefited from WWF-supported snare removal operations, although these efforts have not been as regular or intensive as in the Saola NRs. There are no active patrols in Palé, as it is outside of the Xe Sap NPA.

### Data collection and preparation

We conducted systematic camera-trapping and leech surveys from November 2014 – December 2016. We set up a total of 140 camera-trap stations: 53 stations in Bach Ma NP, 21 in the Hue Saola NR, 25 in the Quang Nam Saola NR, 15 in eastern Xe Sap NPA, and 26 in the Palé area (Fig. 1; Table S2). Stations were spaced approximately 2.5 km apart (mean = 2.47 ± 0.233), aiming at spatial independence of sampling locations, and left in the forest for a minimum of 60 days (mean = 71.60 ± 16.39). Cameras were set 20 – 40 cm off the ground, operational 24 hour/day, and programed to take a three-photo burst with no delay between photographic events. To maximize detection probabilities, we set two camera-traps (Hyperfire Professional PC850, Reconyx^®^, Holmen, USA) at each station facing different directions. We treated the two cameras as a single station in our analyses. Camera-trap data was managed using the package *camtrapR* (Niedballa et al, 2015). We excluded arboreal species from our final species list, as these species are unlikely to be reliably detected by camera-traps placed at ground-level (Abrams et al, 2018). We also removed rodents and squirrels, given the difficulty of identifying these mammals to species-level using camera-trap images alone, and all domestic animals.

We complemented camera-trapping with the collection of terrestrial haematophagous leeches around the camera-trap stations. Leeches were collected once during camera-trap setup and again during retrieval. In Vietnam, leeches were collected in 20 × 20m sampling plots set up to assess microhabitat characteristics (see below). In Laos, we collected leeches in a grid around each camera trap station, with one camera-trap station per grid. We altered the leech collection strategy in Laos because sampling occurred during the dry season; increasing spatial coverage around the stations allowed us to collect leech numbers similar to the Vietnam sites. We separated the two types of leeches, brown and tiger, because the leeches potentially differ in their feeding behavior (Schnell et al 2015). All leeches of the same type from the same station and occasion were combined and processed as one leech bulk sample. Leeches were immediately placed in RNA*later* and stored long-term at −20° C.

Leeches were processed using the laboratory procedures and bioinformatics pipeline described in Axtner et al (2019). The workflow is designed to minimize the risk of false positives that could arise from laboratory artifacts or misidentification during taxonomic assignment. To address these risks, it employs different levels of replication (i.e. extraction, PCRs), a curated reference database, and the probabilistic taxonomic assignment method PROTAX (Somervuo et al, 2017) that has been shown to be robust even when reference databases are incomplete (Rodgers et al., 2017, Richardson et al., 2017). Leech samples were digested, DNA was extracted, and then mitochondrial target DNA of host species was amplified with PCR and sequenced using Illumina high-throughput sequencing. We trained PROTAX models and weighted them toward 127 mammal and bird species expected to occur in the study area by assigning a prior probability of 90% to these species and a 10% probability to all others (Somervuo et al, 2017; see Table S2 for full weighted species list). Our protocol was slightly modified from Axtner et al (2019) in that we amplified the mitochondrial marker 16S in six PCR replicates for all samples and used genetic markers 12S and CytB only for samples where taxonomic assignment was still uncertain due to interspecific invariance or missing references (e.g. porcupines, viverrids, muntjacs). We accepted a species assignment when it was present in at least two independent PCR replicates (Axtner et al. 2019, Abrams et al. 2019). As with the camera-trapping data, we excluded arboreal species, rodents, squirrels, and domestic animals from the final species list.

### Covariates

We hypothesized that mammal occurrence may be influenced by both environmental and anthropogenic factors. We measured three environmental features that characterize different aspects of microhabitat structure: canopy closure, vegetation density, and leaf litter. We used canopy closure as an indication of forest degradation, with lower values representing more disturbed habitat (Chazdon, 2003). Previous studies have shown that vegetation density may be an important microhabitat feature for some tropical mammals (Goulart et al, 2009; Martin et al, 2015; Mathai et al, 2017). Leaf litter impacts multiple aspects of vegetation community composition (Facelli & Pickett, 1991). It is also an important microhabitat for invertebrates and small vertebrates (Burghouts et al, 1992; Vitt & Caldwell, 1994), which are important food resources for insectivores and small carnivores.

To assess microhabitat features, we set up a 20 × 20 m plot around the camera-trap stations, with the centerpoint halfway between the two cameras, and oriented along the cardinal axes. To measure canopy closure we took vertical photographs at the centerpoint and at the corners of the grid. Canopy photographs were manually converted to black and white images using the GNU Image Manipulation Program (GIMP, 2017). We calculated percentage canopy closure (white pixels) for each image using *R 3*.*4*.*0* (R Development Core Team, 2016). Values for each image were averaged to give a single canopy closure value for each station. To measure vegetation density we took photographs in each cardinal direction of a 1 × 1.5 m orange sheet positioned 10 m from the centerpoint. Photographs were processed using the canopy closure protocol, giving a single average vegetation density value for each station. We measured leaf litter percent cover in nine 1 x 1 m subplots located at the centerpoint, 10 m from the centerpoint in each cardinal direction, and at the plot corners. Each subplot was visually assigned a value from 0 to 4 based on the amount of leaf litter versus bare ground visible in each plot. Leaf litter values were averaged to give a single value for each station. For a detailed explanation of the microhabitat assessment see Abrams et al (2018).

In addition to the environmental covariates, we measured anthropogenic features that approximate hunting pressure. We use proxies for hunting pressure, rather than direct measures, for two reasons. First, we are not aware of any existing datasets that directly measure hunting pressure within our study sites. Second, robustly assessing poaching represents a difficult undertaking because illegal hunting is such a cryptic phenomenon. Although some studies have used presence or absence of people from camera-trapping data to represent direct measures of hunting pressure (Dias et al, 2019), such measures are not applicable in our landscape, because some local communities are allowed to legally enter the study sites to collect non-timber forest products. Further complicating the situation is the fact that these local people may engage in both legal non-timber forest product collection and illegal hunting in order to maximize potential profit. Given the difficulties in assessing hunting directly, we used measures of accessibility as proxies for hunting pressure in our study sites. Previous studies have shown accessibility and hunting to be correlated (Rao et al, 2005; Espinosa et al, 2014; Koerner et al, 2017). We used three covariates that capture different aspect of accessibility: distance from major cities, village density, and least cost path from major roads. We used city distance as a proxy for hunting pressure captured at the landscape scale. Although we measure distance to the nearest major city (Hue or Da Nang, both with population > 350,000), we also interpret this covariate as an approximation of accessibility to the densely-populated coastal areas of Vietnam. We chose to measure distance to the cities, rather than other points along the urbanized coastal areas, because Hue and Da Nang are known to be major hubs for the illegal wildlife trade (VanSong, 2003; Sandalj et al, 2016). Given the volume of bushmeat that passes through these markets (Sandalj et al, 2016), it is likely that these urban population centers create substantial natural resource demand shadows across the landscape, as has been shown in other tropical regions (Ape Alliance, 1998). We derived the city distance covariate by calculating the Euclidean distance from the camera-trap stations to the nearest major city using the package *gDistance* (Van Etten, 2017), then taking the lower of the two values. The city distance covariate is measured in meters, with increasing values indicating more remote areas. We then took the log of the covariate to approximate the non-linear effect that increasing distance likely has on accessibility. Village density serves as a proxy for hunting at the local scale. Local villagers often supplement their income by providing bushmeat to the bushmeat markets in regional towns and cities, and are therefore a primary driver of poaching in the central Annamites (MacMillan & Nguyen, 2014). Studies in other tropical regions have demonstrated mammal depletions surrounding local villages (Rao et al, 2005; Koerner et al, 2017; Abrahams et al, 2017). To calculate village density we first created a ground-truthed point shapefile layer documenting local villages around our study sites. We then created a heatmap in QGIS 2.18.9 (QGIS Development Team, 2016) using the village shapefile as the input point layer. To create the heatmap, we used the default quartic kernel decay function and set the radius to 15 km. The village density radius was chosen so that all individual sampling stations in our study landscape were covered in the final heatmap. Observations in the field indicate that all stations, even those in the most in the most remote areas, were subject to some level of hunting pressure. We then used the extract function in the raster package (Hijmans, 2019) to obtain heatmap values for each station. The village density covariate is unitless, with lower values indicating areas that are more remote. Finally, the least cost path covariate also serves as a proxy for hunting pressure at the local scale. However, it differs from the village density measure in two fundamental ways. First, the least cost path covariate explicitly incorporates accessibility based on terrain ruggedness characteristics, therefore providing a more accurate representation of remoteness than linear measures. Second, we calculated the least cost path covariate over three time periods (1994, 2004, and 2014) to better capture the amount of time that an area has been subjected to poaching pressure. The least cost path covariate therefore captures both spatial and temporal dimensions. To create the least cost path covariate we first used the timelapse function in Google Earth Engine (Gorelick et al, 2017) to generate a GIS layer of major roads in and around our study site for three time periods: 1994, 2004, and 2014. We converted the roads layer to points in QGIS. Next, we used the R package *movecost* (Alberti, 2018) to calculate travel time along least cost path routes from the stations to the nearest 50 points along the road layer, using a shuttle radar topography mission (SRTM) 30 m digital elevation model as the cost surface raster, and then selected the lowest value as the final least cost path value. We averaged the three values to give a single least cost path value for each station, which we use as an approximation of the time that an area has been accessible over the past 20 years. The roads least cost path covariate is measured in hours, with higher values indicating areas that take longer to access, and are therefore more remote.

We also included elevation as a covariate in our models. We consider elevation as both an anthropogenic and ecological covariate. Because higher elevation areas are more difficult to access, elevation serves as a measure of remoteness within our landscape. Elevation is also linked to a complex range of ecological attributes in the central Annamites, including subtle variations in forest structure and microclimate (Tordoff et al, 2003; Long, 2005).

We standardized all covariates. We tested for correlations between all possible pairs of covariates using Pearon’s correlation plots. None of our covariates were highly correlated (r < 0.6; Fig. S2).

### Modeling framework

We adopted a hierarchical multi-species occupancy model to estimate species occupancy and richness (Dorazio & Royle, 2005; Dorazio et al., 2006). Occupancy models estimate the probability of species occupancy, *ψ*, while accounting for species detection, *p*, using repeated species detection/non-detection data collected across multiple sampling locations (MacKenzie et al., 2003). To convert camera-trapping data to an occupancy format, we divided the active camera-trapping time for each station into 10-day sampling periods, yielding a minimum of six occasions for each station. We chose to use a 10-day sampling period to minimize zero-inflation in the detection history matrix. We treated each leech collection event as a separate occasion for the stations. We defined *z*_*ij*_ as the true occupancy state (0 or 1) of species *i* at sampling station *j*. Occupancy state can be modeled as a Bernoulli random variable with the success probability *ψ*_*ij*_, the occupancy probability of species *i* at site *j*. We defined *p*_*ijk*_ as detection probability for species *i* at station *j* during the *k*th sampling occasion, and *y*_*ijk*_ the observation (i.e., *y*_*ijk*_ = 1 if species *i* is observed at site *j*, occasion *k*, and 0 otherwise). Observing a species is conditional on its occurrence, so that *y*_*ijk*_ can be modeled as a Bernoulli random variable with success probability *z*_*ij*_ · *p*_*ijk*_.

Covariate effects on both parameters can be modeled on the logit scale. We included habitat and anthropogenic covariates on *ψ*_*ij*_ to investigate their potential effects on species occurrence. To avoid overparametizing the model, we first ran single-covariate models using each of the seven covariates that we selected *a priori*, and assessed covariate importance by evaluating effect sizes for each species in the community. Because the environmental covariates did not show strong effects on occupancy, with all species having 95% BCIs overlapping zero and most species showing overlapping 75% Bayesian Confidence Intervals (BCIs) (Fig. S3), these covariates were not included in the final model. Our final community model included four covariates on *ψ*_*ij*_: city distance, village density, roads least cost path, and elevation. Following Abrams et al (2019), we used survey method (camera-trap, brown leech, or tiger leech) as a covariate on *p*. We accounted for varying survey effort by including number of days each camera-trap station was operational during each 10-day occasion or number of leeches per sample on *p*.

We implemented the models in a Bayesian framework using JAGS (Plummer, 2003) accessed through the package rjags (Plummer, 2018). We used vague priors (e.g. normal distributions with mean zero and variance 100 for community-level occupancy and detection coefficients). We ran three parallel Markov chains with 250,000 iterations, of which we discarded 50,000 as burn-in. We assessed chain convergence using the Gelman-Rubin statistic, with values close to 1 indicating convergence (Gelman et al, 2004). We report results as posterior mean and standard deviation. We consider a coefficient to have strong support if the 95% Bayesian confidence interval (95% BCI, the 2.5% and 97.5% percentiles of the posterior distribution) does not overlap zero, and moderate support if the posterior 75% BCI does not overlap zero. The full model description is provided in Appendix 1.

To test for spatial autocorrelation in response variables not accounted for by predictor variables, we followed an approach put forth by Moore and Swihart (2003). We calculated Moran’s I using the moransI function from the R package lctools for residuals from occupancy models using a neighborhood distance of 2.5 km (average spacing of our sampling stations). We only found evidence of low to moderate spatial autocorrelation in occupancy model residuals in only 2 of the 23 species analyzed. We acknowledge that for these species we may underestimate occupancy. However, our analysis is concerned with comparisons of patterns across study sites, not among species, and for a given species, any bias should be similar across the sites. Estimates of Moran’s I and associated p-values for species are shown in Appendix 2.

To predict species richness across the landscape we first divided the study area into 200 × 200 m grid cells. We included proposed extensions for the Hue and Quang Nam Saola Nature Reserves in the prediction area. Next, we derived covariate values for each cell. For the city distance, village density, and roads least cost path covariates we followed the same protocols described above. Elevation values were extracted from an SRTM 30 m digital elevation model (see Fig. S4 for covariate rasters). We used estimates of the coefficients from the multi-species model linking covariates to occupancy probability to predict occupancy values for each species and grid cell and then summed the occupancy probabilities for all species per cell to produce species richness maps. To highlight areas of high richness for conservation-priority species, we produced a separate species richness map for the endemic species and those listed as Near Threatened or higher on *The IUCN Red List of Threatened Species*. To provide a further level of detail for the endemic species, we also produced single-species occupancy maps for Annamite striped rabbit Annamite dark muntjac and Owston’s civet. We note that, although our sampling stations only covered part of the study sites, covariate values at the stations were largely representative of values across the sites. When we filtered the raster cells to remove cells that fell outside the range of our covariates at the sampling stations, we found that a small number raster cells were excluded. However, we present the full prediction maps here, both because the differences between the complete and filtered rasters were minor, and for visualization purposes (see Fig S5 for modified prediction maps).

We further used a modified Bray-Curtis index to assess compositional dissimilarity among the five study sites. The Bray-Curtis index calculates dissimilarity values by comparing composition in a reference assemblage with one or more target assemblages (Bray & Curtis, 1957). We adapted the index to compare predicted species occupancy probabilities between all possible site combinations, following the general framework proposed by Giocomini & Galetti (2013). To do this we sampled random values from the posterior distributions of species-specific occupancy probabilities for both the focal and target study sites. We repeated this procedure 30,000 times using Monte Carlo sampling to generate a distribution of values and took the mean of the posterior distribution. The final value gives an indication of how dissimilar the predicted community-level occupancies are among the sites. Dissimilarity values can range between −1 and 1. A value of 0 indicates no differences in occupancy between the focal and reference sites, a value of 1 indicates complete dissimilarity with the reference site having higher occupancies than the focal site, and a value of −1 indicates complete dissimilarity with the focal site having higher occupancies than the reference site. We calculated Bray-Curtis dissimilarities first for the entire community, and then for endemic and threatened species. Further details on the Bray-Curtis dissimilarity index are provided in Appendix 3.

## Results

We obtained data from 139 camera-trap stations totaling 17,393 trap-nights (Table S1). The camera-trapping yielded 5 261 independent detection (Δ = 60 min between subsequent pictures of the same species at the same camera trap) of 27 terrestrial mammals. We identified all mammals to species, with the exception of the ferret badgers (*Melogale personata* and *M. moschata*) and pangolins (*Manis pentadactyl* and *M. javanica*), which we identified to the genus level due to the difficulty of identifying to species using camera-trap photographs, and the Annamite dark muntjac species complex *Muntiacus rooseveltorum / truongsonensis*, due to its unresolved taxonomic status. Our final species list resulted in 22 mammals. We obtained 193 leech samples totaling 2,043 leeches (1,888 brown, 155 tiger) from 98 stations (mean leeches / station = 21, standard deviation = 22; Table S1). We were able to amplify and sequence DNA from 104 samples. PROTAX identified 25 mammals to the species level and 7 to the genus level. The final species list from the leeches included 19 terrestrial mammals. Overall, the two survey methods provided similar species lists. The exceptions were pangolin, pig-tailed macaque *Macaca leonina*, spotted linsang *Prionodon pardicolor*, and yellow-bellied weasel *Mustela* kathiah, which were detected only in the camera-traps, and marbled cat *Pardofelis marmorata*, which was detected only in the leeches. The final species list used for the community occupancy analysis included 23 mammals. Four of these were threatened, three were Annamite endemics, and one species fit both categories. The full species list and classifications can be found in Table S3.

Detection probabilities (*p*) within the mammal community varied among species and with respect to survey method. Estimates of occupancy (*ψ*) showed extreme heterogeneity among individual species. Estimates of *p* and *ψ* can be found in Fig. S6, and full model results are provided in Table S4.

Species-specific responses to the covariates were highly variable within the community (Fig. 2). Six species showed a moderate positive relationship with city distance, three species showed a moderate negative relationship, and four species showed a strong negative relationship. There was a moderate positive effect of elevation on occupancy for nine species and a strong positive effect for four species. One species showed a moderate negative response to elevation and three species showed a strong negative response. There was a moderate positive relationship for elevation at the community level. Five species showed a moderate positive response to the road least cost path covariate. Three species showed a moderate negative relationship with this covariate and one species had a strong negative response. Most species showed a negative relationship with village density, with 16 species having a moderate negative relationship and four species showing a strong negative relationship. The community response for the village density covariate showed a strong negative response.

**Figure 2:**
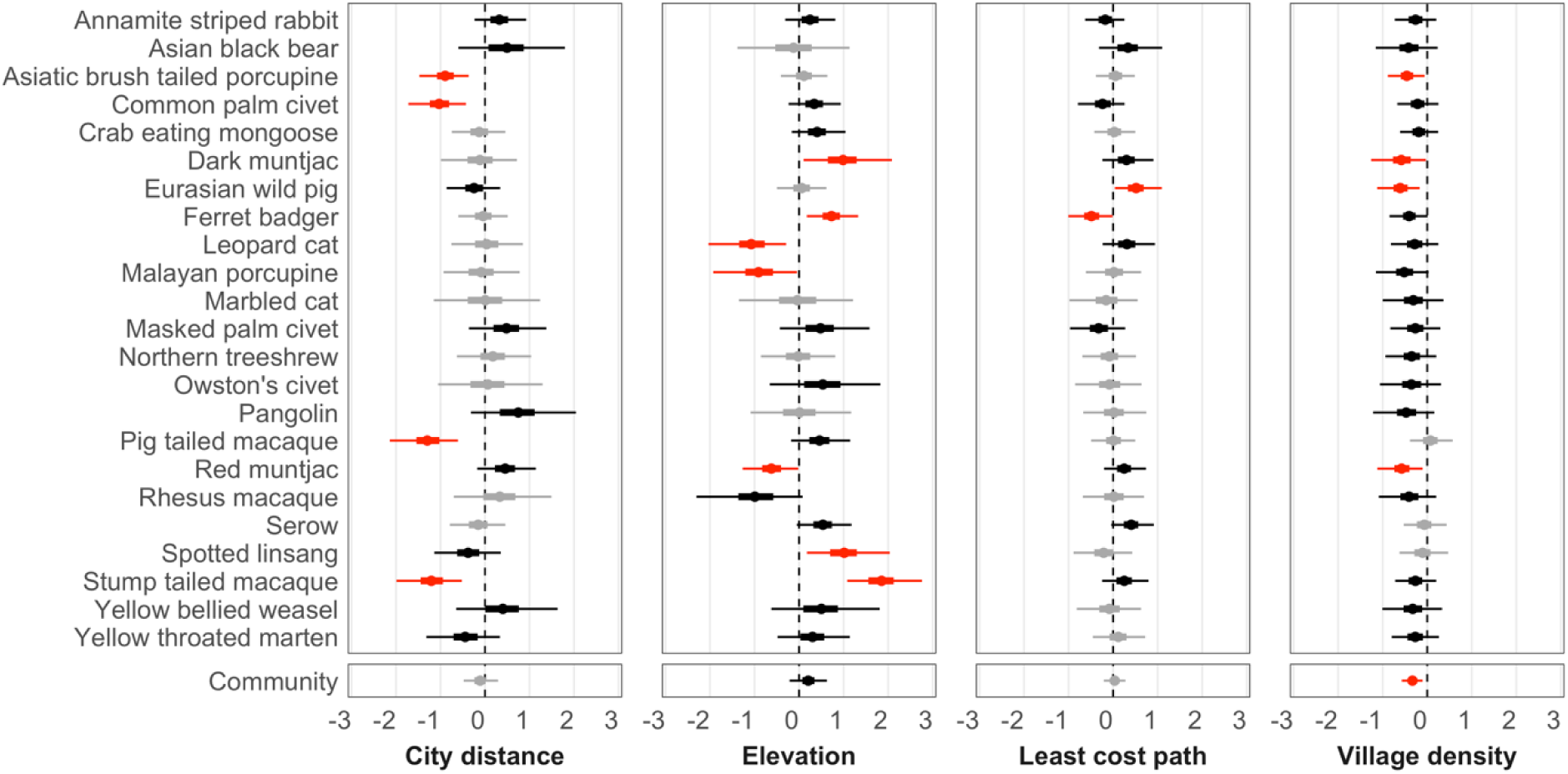
Standardized beta coefficients (mean, 95% BCI, 75% BCI, on the logit scale) showing covariate effects on species occupancy Gray bars show relationships in which the 75% Bayesian Credible Interval (BCI) does not overlap zero, black bars indicate that the 75% interval does not overlap zero but the 95% interval does overlap zero, and red bars indicate that the 95% interval does not overlap zero. The community response is shown in the lower panel.

Average predicted species richness for the full community was substantially lower than the total number of species detected in the study, and was similar among the study sites (Table 1, Fig. 3A). The Palé area had slightly higher predicted species richness than the other four areas. Richness of threatened and endemic species followed a similar pattern, with all sites showing low richness relative to the total number of conservation-priority species detected, and the highest richness in the Palé area (Table 1, Fig. 3B). Predicted occupancies for the three Annamite endemic mammals showed heterogeneity among species and sites (Table 1, Fig. 4). Annamite dark muntjac had the highest predicted occupancy, followed by Annamite striped rabbit, followed by Owston’s civet. All three endemics had highest predicted occupancies in the Palé area, followed by Xe Sap NPA, Quang Nam Saola NR, Hue Saola NR, and finally Bach Ma NP. Single-species occupancy predictions for all species can be found in S7.

**Table 1:** Predicted species richness and species occupancies (mean ± SD) for five study sites in the central Annamites landscape, from multi-species community occupancy model fit to 23 mammal species. Full community indicates richness for all 23 species. Threatened and endemic species indicates richness for 10 species that are endemic and / or listed as Near Threatened or higher on *The IUCN Red List of Threatened Species*. Species occupancy shows predicted occupancies for each of the three Annamite endemic mammals. Occupancy values range from 0 to 1.

|  |  | Study site |  |  |  |  | All sites |
| --- | --- | --- | --- | --- | --- | --- | --- |
|  |  | Bach Ma NP | Quang Nam Saola NR | Hue Saola NR | Xe Sap NPA | Palé |  |
| Species richness | Full community | 6.52 ± 1.39 | 6.32 ± 1.00 | 7.00 ± 1.24 | 6.89 ± 0.98 | 8.25 ± 1.13 | 7.02 ± 1.41 |
|  | Threatened and endemic species | 1.79 ± 0.77 | 1.80 ± 0.65 | 1.85 ± 0.66 | 2.10 ± 0.70 | 2.97 ± 0.85 | 2.12 ± 0.89 |
| Species occupancy | Annamite striped rabbit | 0.15 ± 0.04 | 0.24 ± 0.04 | 0.20 ± 0.05 | 0.28 ± 0.03 | 0.36 ± 0.05 | 0.24 ± 0.09 |
|  | Annamite dark muntjac | 0.24 ± 0.18 | 0.35 ± 0.15 | 0.30 ± 0.18 | 0.41 ± 0.17 | 0.66 ± 0.19 | 0.39 ± 0.24 |
|  | Owston's civet | 0.05 ± 0.02 | 0.06 ± 0.02 | 0.05 ± 0.02 | 0.08 ± 0.02 | 0.13 ± 0.05 | 0.07 ± 0.05 |

**Figure 3:**
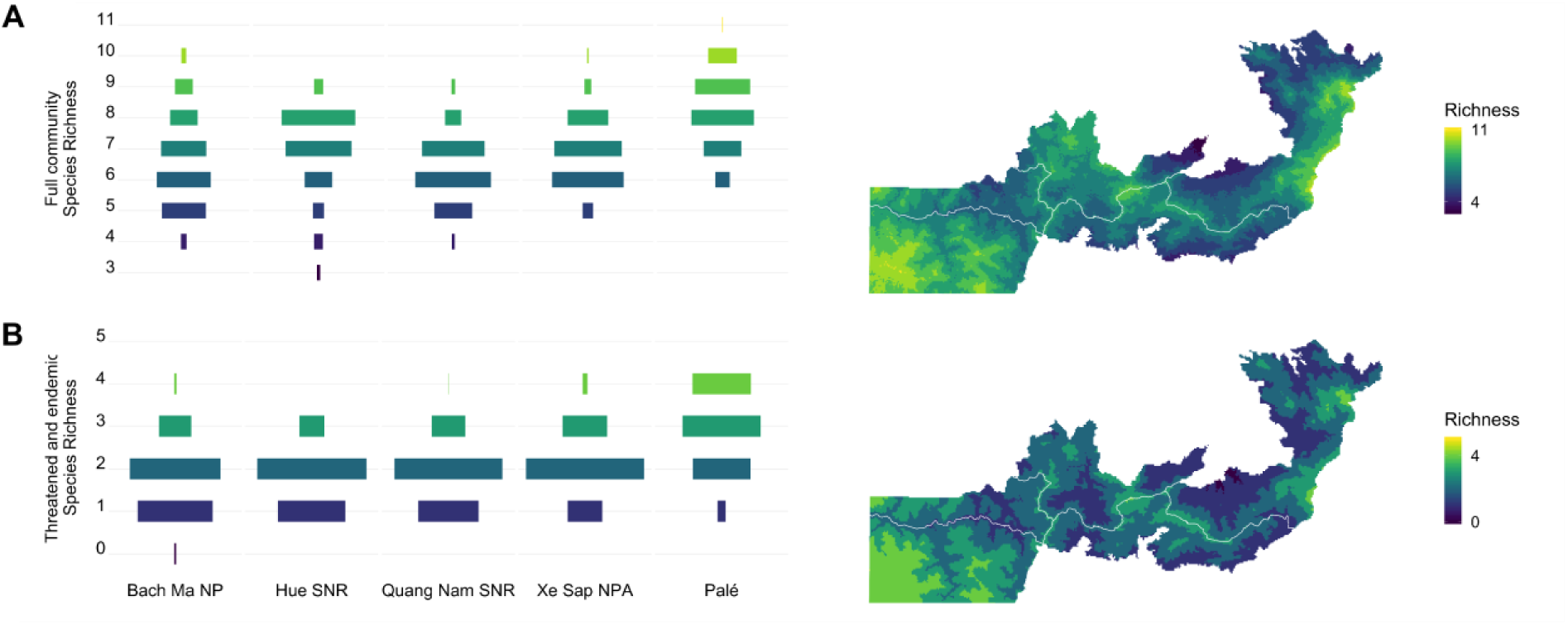
Predicted species richness across five study sites in the central Annamites landscape with histogram showing proportion of cells in each study area for predicted species numbers (left panel) and prediction map (right panel) based on community occupancy model fit to camera-trapping and iDNA data for 23 mammal species. (A) Predicted richness for all species. (B) Predicted species richness for threatened and endemic species.

**Figure 4:**
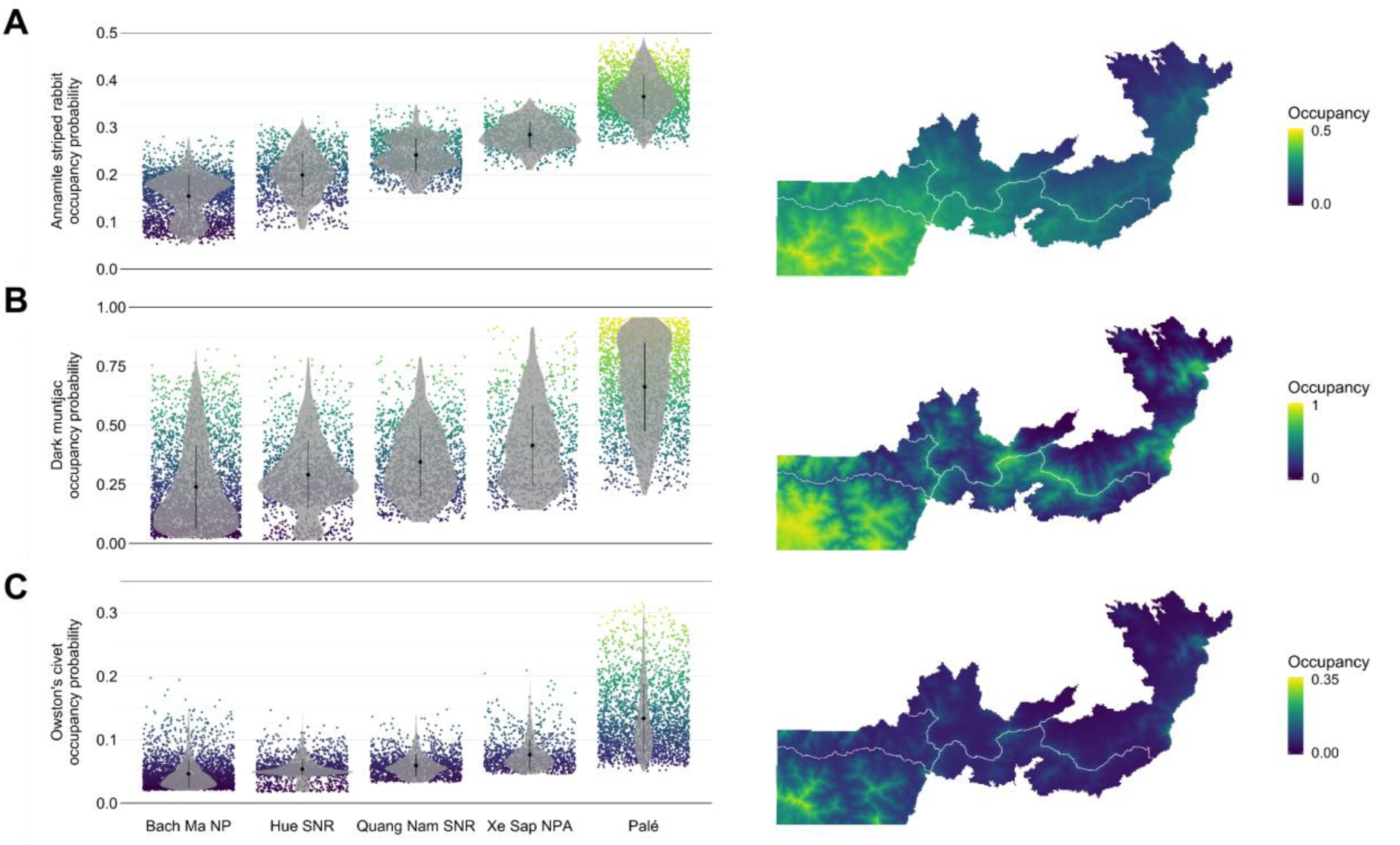
Predicted occupancies for three Annamite endemic species with violin plot showing predicted occupancy values for each of the five study sites (left panel) and prediction map (right panel) based on community occupancy model fit to camera-trapping and iDNA data for 23 mammal species. (A) Predicted occupancy for Annamite striped rabbit. (B) Predicted occupancy for Annamite dark muntjac. (C) Predicted occupancy for Owston’s civet. Note that, for visualization purposes, occupancy values are scaled independently for each species. Single-species prediction maps with standardized scaling for occupancy values can be found in Fig. S7.

The Bray-Curtis dissimilarity index values showed similar values for full community occupancies among Bach Ma NP, the Hue and Quang Nam Saola NRs, and Xe Sap NPA (Table 2). However, the Palé area had negative and high defaunation index values when compared to every other site, indicating that occupancies for the full suite of species are higher for this area. Dissimilarity values for endemic and threatened species showed a similar pattern. The fact that Palé area showed negative and higher dissimilarity values for both the full community and for conservation-priority species suggests that, within the context of species occurrence, the site has undergone less severe defaunation than the other sites.

**Table 2:** Bray-Curtis dissimilarity values (mean ± SE) calculated using predicted occupancy values per site for all mammal species (top value) and for threatened and endemic species (bottom value). Values range from 0 to 1, with 0 indicating complete similarity in species occupancies between the sites, and 1 indicating complete dissimilarity.

|  | Bach Ma NP | Quang Nam Saola NR | Hue Saola NR | Xe Sap NPA | Palé |
| --- | --- | --- | --- | --- | --- |
| Bach Ma NP | 0 | -0.0006 ± 0.0205<br>0.0431 ± 0.0205 | 0.0406 ± 0.0119<br>0.0049 ± 0.0119 | 0.0350 ± 0.0233<br>0.0841 ± 0.0233 | 0.1338 ± 0.0315<br>0.2547 ± 0.0315 |
| Quang Nam Saola NR | 0.0006 ± 0.0205<br>-0.0431 ± 0.0205 | 0 | 0.0412 ± 0.0206<br>-0.0382 ± 0.0205 | 0.0357 ± 0.0061<br>0.04127 ± 0.006 | 0.1346 ± 0.0193<br>0.2150 ± 0.0193 |
| Hue Saola NR | -0.0406 ± 0.0119<br>-0.0049 ± 0.0119 | -0.0412 ± 0.0205<br>0.0382 ± 0.0206 | 0 | -0.0055 ± 0.0206<br>0.0793 ± 0.0206 | 0.0938 ± 0.0297<br>0.2503 ± 0.0297 |
| Xe Sap NPA | -0.0350 ± 0.0233<br>-0.0840 ± 0.0233 | -0.0357 ± 0.0061<br>-0.0413 ± 0.006 | 0.0055 ± 0.0206<br>-0.0792 ± 0.0206 | 0 | 0.0995 ± 0.0174<br>0.1754 ± 0.0174 |
| Palé | -0.1338 ± 0.0315<br>-0.2547 ± 0.0315 | -0.1346 ± 0.0193<br>-0.2149 ± 0.0192 | -0.0938 ± 0.0297<br>-0.2502 ± 0.0297 | -0.0995 ± 0.0174<br>-0.1754 ± 0.0174 | 0 |

## Discussion

Our study highlights the landscape-scale effects of unsustainable hunting on the occurrence and distribution of terrestrial mammals within a tropical biodiversity hotspot. For a structurally-intact tropical rainforest habitat, average predicted species richness (7.02) was low (Table 1; Fig. 3A & B). For comparison, Deere et al (2018) found a predicted species richness of 14.12 in logged forests in Malaysian Borneo, and a richness of 4.54 in adjacent oil palm plantations. Given the largely homogenous landscape-scale forest structure and habitat in our study sites, the low predicted species richness is likely indicative of a community that has undergone severe hunting-driven defaunation. The extent of faunal impoverishment is further supported by the fact that we failed to record almost half of the mammal community would be expected to occur in these sites based on historical distribution maps (Tilker & Abrams, *in press*).

At the landscape level predicted richness was broadly similar among the five study sites, although the more remote Palé area showed the highest richness values, especially for the endemic and threatened species (Fig. 3). The Bray-Curtis dissimilarity values show a similar geographic pattern, with the Palé area having higher dissimilarity values compared to the other four areas, indicating less defaunation (Table 2). For the three endemic species, Bach Ma NP showing the lowest predicted occupancy, followed by the Hue Saola NR, Quang Nam Saola NR, Xe Sap NPA, and Palé. These findings indicate a strong landscape-scale defaunation gradient for the three endemic species (Fig. 4). Our covariate responses suggest that this gradient reflects an increasing level of remoteness (Fig. 2). Bach Ma NP lies near densely-populated coastal areas of Vietnam, has lower average elevations, and has been accessible for decades by a well-established road network. In the westernmost section is the Palé area, which is far from major cities, has few villages, higher elevations, and has only recently been accessible by road. The Saola NRs and Xe Sap NPA fall between these two extremes. Furthermore, these areas have had some level of active enforcement in the last few years, which may have slowed the decline of mammal populations.

Our results provide information that is directly applicable to conservation planning in this landscape. From a biogeographic perspective, protecting the Palé area is a top priority for the conservation of threatened and endemic species. Indeed, it may be the only place in our survey sites to harbor Owston’s civet Asian black bear *Ursus thibetanus*, and marbled cat (Fig. S7). Our predictive maps offer a robust scientific framework to support ongoing initiatives to grant Palé formal protected area status as a first step to implementing active protection measures. Our maps also provide information to guide targeted snare-removal efforts within protected areas. This information is especially useful for the Hue and Quang Nam Saola NRs, where WWF and local partners are operating snare-removal teams, but have not yet been able to significantly reduce snaring pressure across the wider protected area complex (Wilkinson, 2016). We suggest that, to maximize the impact of snare-removal efforts on conservation-priority species, the teams should focus on the more remote areas along the border of the two reserves, and in the border area of the Quang Nam Saola NR and Bach Ma NP. It is possible that these areas have maintained higher occupancies of conservation-priority species because they are more difficult to access and, as a result, have cumulatively experienced less snaring pressure. However, remoteness will not protect these areas for long. An increase in road development in recent years has created a situation where even the most remote locations in the Saola NRs can now be reached within a single day from the nearest access point, meaning that no area is inaccessible for a motivated hunter. Given the likely relationship between accessibility and increased hunting pressure, it seems inevitable that, in the absence of scaled-up enforcement efforts, snaring pressure will continue to increase in the more remote areas, especially as other parts of the protected areas become increasingly empty and poachers are forced to travel further distances to maintain comparable levels of offtake (Kümpel et al, 2010). Although threatened and endemic species appear to be absent from much of Bach Ma NP, there are isolated high-elevation areas that should be considered for intensive anti-poaching efforts. Our models indicate that the border areas of eastern Xe Sap NPA Area have undergone moderate to severe defaunation. Given that this area appears to be heavily hunted by Vietnamese poachers (Tilker, 2014), such a finding is not unexpected. The eastern section of the protected area is nonetheless a top priority for continued protection efforts, both because it may be a stronghold for the endemic Annamite striped rabbit (Fig. S7), and more generally because effective enforcement can serve as a buffer from further cross-border incursions into the Palé area. In a best-case scenario, reducing snaring pressure in core areas within this landscape could not only prevent the local extirpation of conservation-priority species, but also allow their populations to rebound (see Steinmetz et al, 2010 for a case study on large mammal population recovery following mitigation of unsustainable hunting pressure).

In many ways, our study landscape exemplifies classical “empty forest syndrome” (Redford, 1992). All large and medium-sized predators (with the exception of Asiatic black bear), as well as all megaherbivores, appear to be locally extirpated (Tilker & Abrams et al, *in press*). Large ungulates have been hunted out from most of the landscape (Fig. S7). Yet our findings show that even in this empty forest, conservation-priority species still persist, albeit at extremely low occupancies. Based on these results, we suggest that the conservation potential of defaunated landscapes should not automatically be dismissed in the absence of comprehensive surveys. It is important that such surveys use sufficient sampling effort and be conducted over a large spatial extent for two reasons. First, species often show extreme spatial heterogeneity in defaunated landscapes because local extinctions necessarily result in reduced, often patchy, distributions. Surveys over wider areas are more likely to detect remnant populations. Second, working over larger spatial scales may better capture the underlying factors influencing species distribution, which can be especially important in landscapes characterized by complex anthropogenic pressures operating at multiple spatial scales. In our study, it was only by sampling the wider forest complex that we were able to adequately characterize the full spectrum of anthropogenic factors that appear to impact species occurrence patterns. Large-scale surveys require substantial resources. We acknowledge that, with multiple competing conservation objectives and finite resources, landscape-scale surveys may not always be possible. However, we note that because this approach can enhance the efficiency of targeted interventions, it is possible that limited conservation resources may be saved in the long term.

To overcome the challenge of detecting rare and elusive tropical mammal species, we used two complimentary survey methods: camera-trapping and leech collection. Although camera-trapping detected more species overall, leeches provided our sole detection for marbled cat, and doubled the number of records for two rare species, Owston’s civet and Asian black bear. Moreover while camera-trapping detection probabilities were higher for most species in our analysis, leeches had higher average detection rates for both Asian black bear and the endemic dark muntjac (Fig. S7). Our results are consistent with the findings of Abrams et al (2019) that demonstrate the advantages of using both camera-trapping and eDNA to increase detection probabilities for tropical mammals. We further suggest that because utilizing multiple methods may increase detections of rare species, this approach could be especially important when surveying faunally impoverished systems. Future surveys using joint detection methods need not rely only on leeches but could use other sources of eDNA, such as water (Ushio et al, 2017) or ticks (Gariepy et al, 2012), or incorporate other noninvasive sampling techniques, such as acoustic monitoring devices (Kalan et al, 2015). The Bayesian modeling approach that we used, adapted from Abrams et al (2019), is flexible with regard to the underlying detection method used to generate spatial or temporal replicates.

We found that species occurrence in our study area appears to be primarily driven by anthropogenic factors, with no strong influence from the habitat covariates that we assessed in our models (Fig. S3). The lack of a strong signal with the habitat covariates was unexpected, given the importance of vegetation structure in explaining mammal occurrence patterns in other tropical rainforests (Goulart et al, 2009; Mathai et al, 2017; Sollmann et al, 2017). One possible explanation is that, as anthropogenic pressures in a landscape increase, ecological relationships weaken. Several hypothetical scenarios could give rise to this situation. Spatially nonrandom hunting pressure could, for example, differentially impact areas of preferred habitat, leaving higher occupancies in less suitable areas. Alternatively, intensive hunting across a landscape could drive stochastic local extinctions, leaving remnant populations that are distributed randomly with respect to habitat. Regardless of the underlying process, the failure of habitat-based indices to reflect faunal biodiversity, thus the “environmental decoupling” of species-habitat relationships, has broad implications. Biodiversity assessments that rely solely on remote-sensed habitat-based measures may provide information that is inaccurate because they do not accurately capture species occurrence patterns. Recently, a growing number of scientists have called for the development of standardized remote-sensing parameters, often referred to as Satellite Remote-Sensing Essential Biodiversity Variables (SRS-EBVs), to monitor biodiversity at the global scale (O’Connor et al, 2015; Skidmore et al, 2015; Pettorelli et al, 2016). While we acknowledge the value of earth observation data to provide insight into biodiversity patterns and processes at large scales (Bush et al, 2017), our results indicate that remote-sensed habitat-based measures may provide little information on the status or distribution of wildlife in defaunated landscapes. In tropical rainforests subject to hunting pressure, there is likely no substitute for large-scale *in situ* surveys to collect primary biodiversity data.

Our results underscore the importance of incorporating anthropogenic factors in studies that seek to explain or predict species occurrence in landscapes characterized by high human pressure. Furthermore, our findings suggest that to build robust distribution models it may be beneficial to incorporate a diverse suite of anthropogenic covariates that capture different aspects of this pressure. We used measures of village density and city distance as proxies for accessibility at the local and landscape scales, respectively. Previous studies have shown the impact of similar accessibility measures on wildlife communities at different spatial scales (Schuette et al, 2013; Koerner et al, 2017; Torres et al, 2018). Our least cost path covariate adds an additional dimension to these accessibility measures, both because it takes into account the ruggedness of the terrain in our landscape, and because it is calculated over a 20-year window. We see potential for further development of anthropogenic covariates that include both spatial and temporal components. Finally, we use elevation as a proxy for both local accessibility and a complex set of ecological attributes. The relative contribution of anthropogenic and environmental traits to species occurrence along elevational gradients in the Annamites represents an intriguing question. Future studies in the region that measure a wider range of microhabitat characteristics, and ideally are conducted in areas under less severe hunting pressure, may provide insight into this issue.

Given the current magnitude of hunting across the world’s tropical rainforests (Harrison, 2011; Ripple et al, 2016; Benítez-López et al, 2017), and future projections for population growth (Gerland et al, 2014) and road expansion in developing countries (Lawrence et al, 2014), it is likely that defaunation will become increasingly prevalent in tropical regions. Confronting the pantropical defaunation crisis will require a well-resourced, multi-faceted approach from conservation stakeholders worldwide. Because specific threats and potential solutions necessarily depend on local context, effective strategies to prevent unsustainable hunting must be site-specific. One constant that is applicable to conservation initiatives in all tropical hotspots, however, is that resources are limited. We show that, within this context, understanding spatial patterns of defaunation can help stakeholders prioritize areas for conservation activities, and therefore more effectively use finite conservation resources.

