## Supplementary Information for "Identifying conservation priorities in a defaunated tropical biodiversity hotspot"

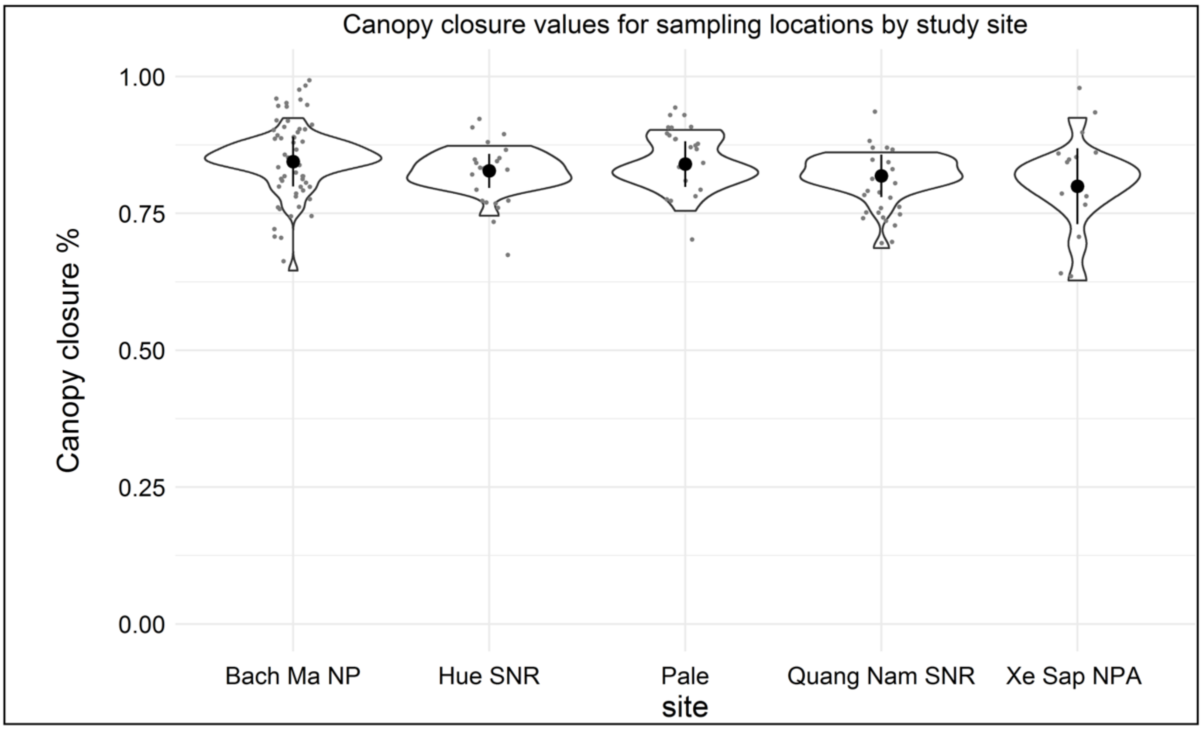


**Figure S1:** Canopy closure values for each study site, assessed using data collected *in situ* around each sampling station. Canopy closure is similar across the study sites, indicating little heterogeneity in forest structure or habitat type. See Abrams et al (2018) for detailed information on *in situ* sampling of canopy closure and processing of canopy photographs.

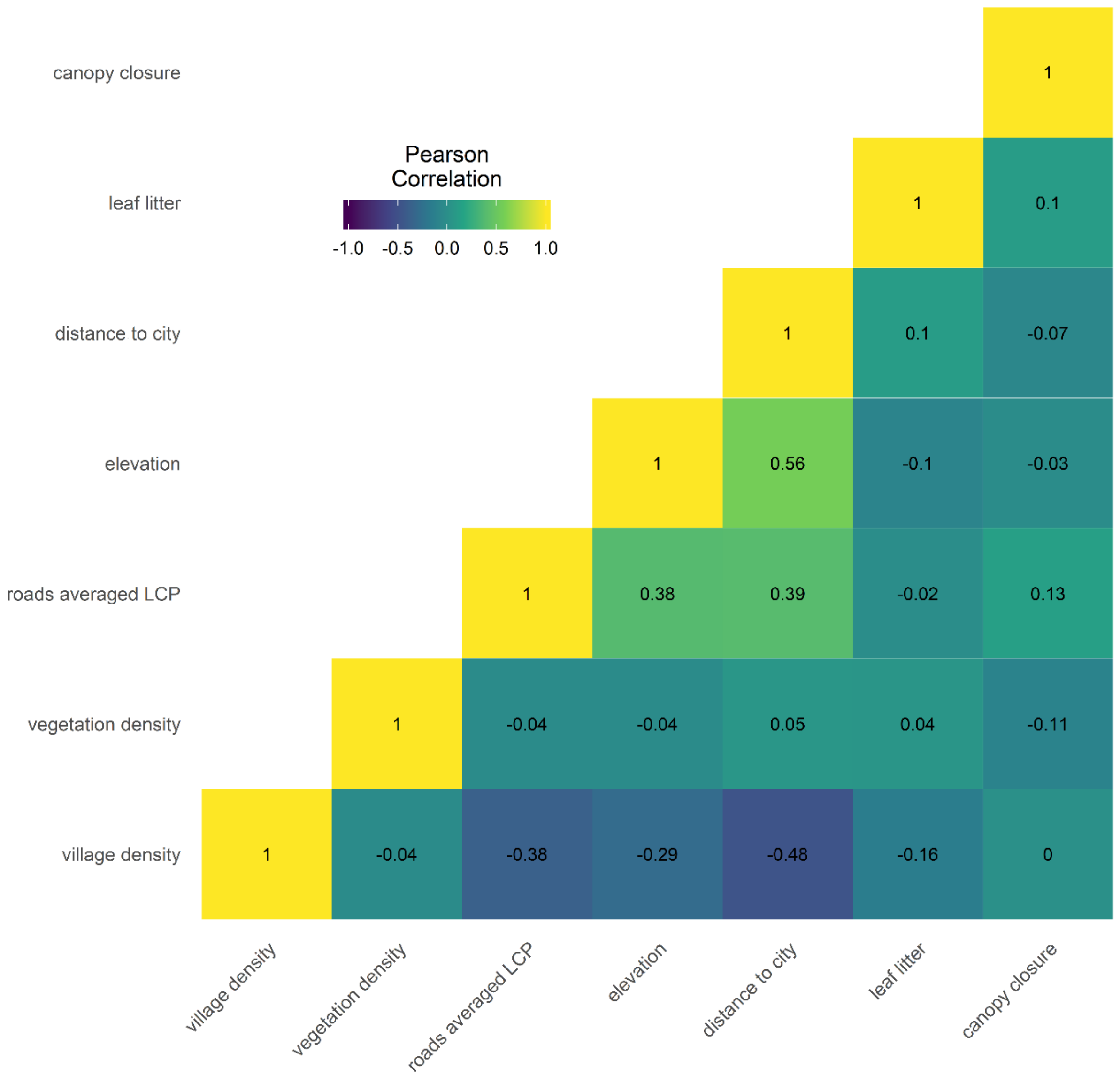


**Figure S2:** Heatmap correlation matrix showing Pearson correlation values for all environmental and anthropogenic covariates.

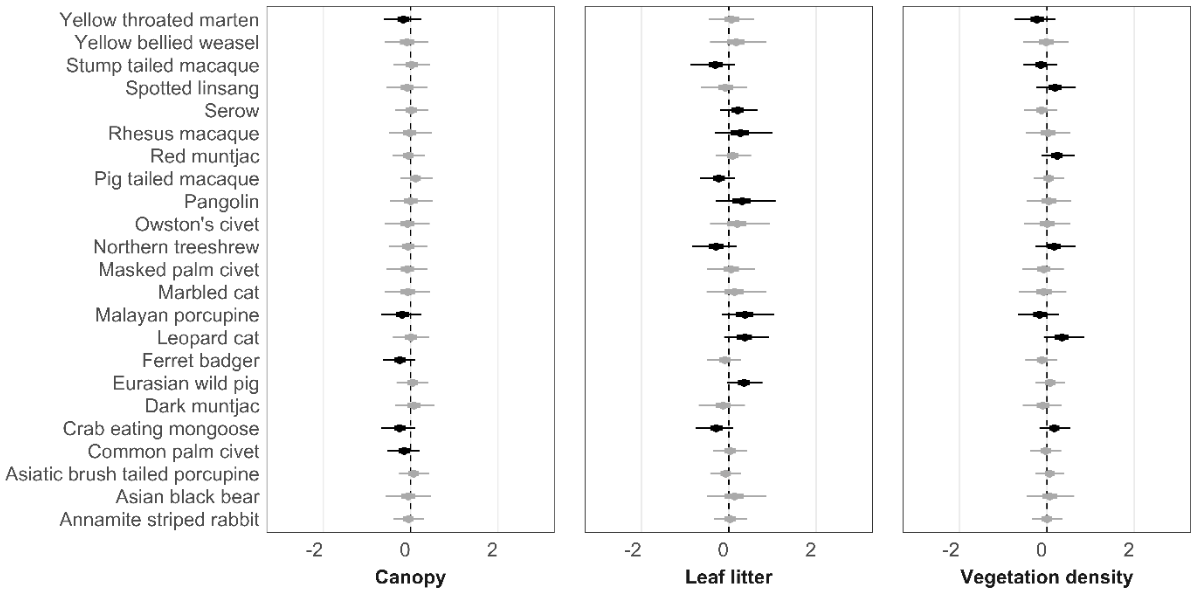


**Figure S3:** Standardized beta coefficients (mean and 95% BCI, on the logit scale) showing environmental covariate effects on occupancy for all species. Gray bars show relationships in which the 75% BCI does not overlap zero, black bars indicate that the 75% interval does not overlap zero but the 95% interval does. No coefficients have 95% intervals that do not overlap zero (see Fig. 2 for comparison).

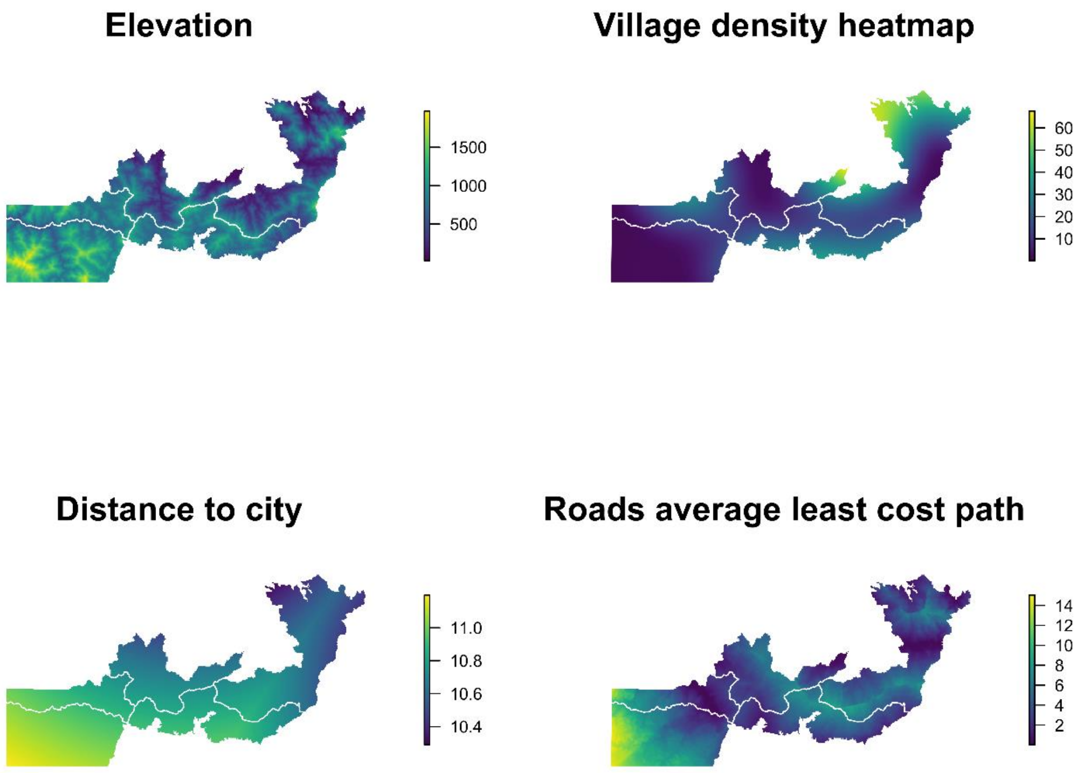


**Figure S4:** Map of four covariates used to make prediction maps. Study sites are demarcated with white lines.

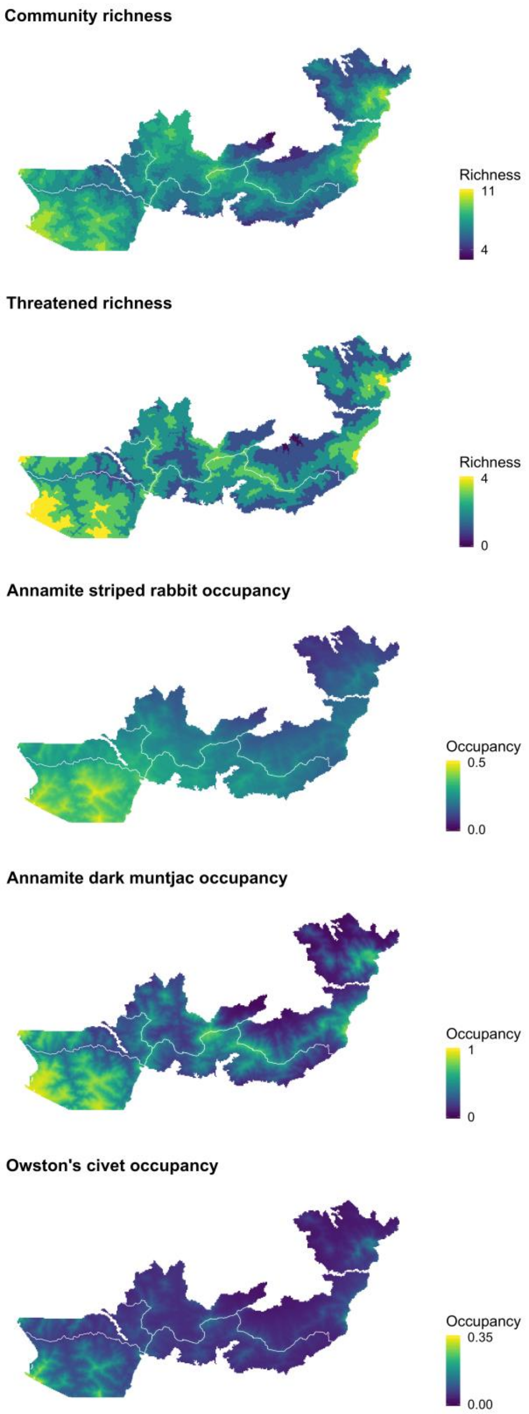


**Figure S5:** Predicted species richness and species occupancy maps with raster cells removed that fall outside the range of covariates at the sampling stations. One extreme lowland area is removed in Bach Ma NP, and one lowland area in eastern Xe Sap NPA, and a remote area that falls to the west of our sampling stations that covers both Xe Sap NPA and the Palé area.

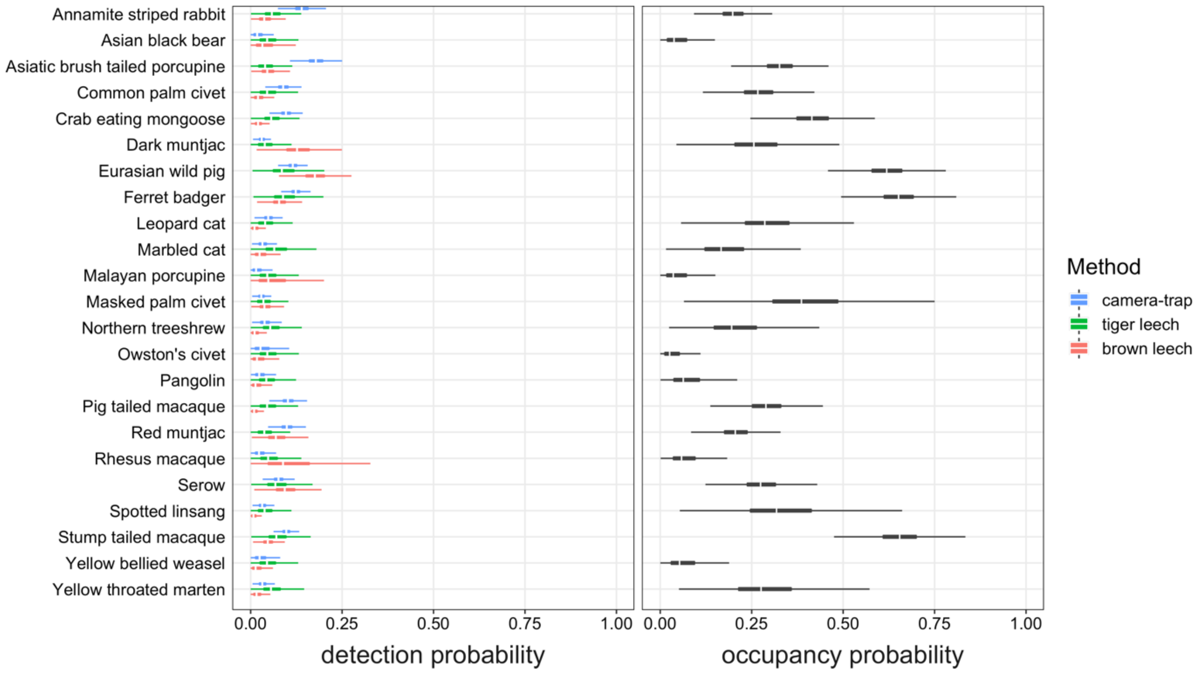


**Figure S6:** Estimated detection probabilities ($p$) for each species and survey method (camera-trap, brown leech, tiger leech; left panel). Estimated occupancy probabilities ($\psi$) for each species (right panel).

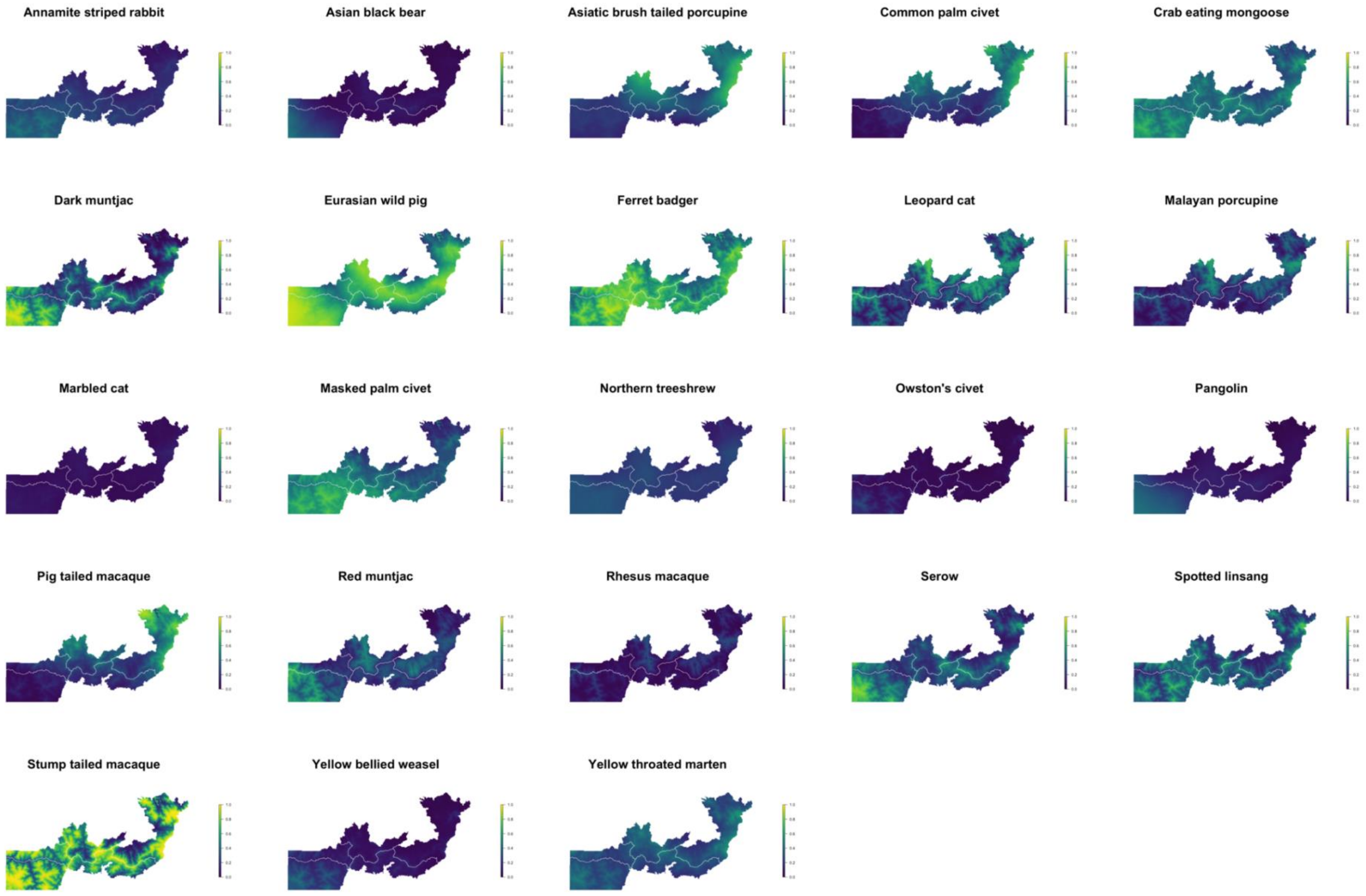


**Figure S7:** Single-species occupancy prediction maps from Bayesian multi-species occupancy analysis fit to 23 mammal species.

| Site | Survey dates | No. stations | No. camera-trap nights | No. leeches / samples |
|---|---|---|---|---|
| Bach Ma National Park | Nov. 2014 - Jan. 2015 | 53 | 6,805 | 201 / 39 |
| Hue Saola Nature Reserve | Aug. - Dec. 2015 | 21 | 2,600 | 292 / 34 |
| Quang Nam Saola Nature Reserve | Aug. - Dec. 2015 | 25 | 2,670 | 641 / 59 |
| Xe Sap National Protected Area | July - Dec. 2016 | 15 | 1,634 | 176 / 17 |
| Ban Palé | July - Dec. 2016 | 25 | 3,684 | 773 / 44 |
| **Total** | Nov. 2014 - Dec. 2016 | **139** | **17,393** | **2,043 / 193** |

**Table S1:** Information for each study site showing survey periods, number of sampling stations, camera-trapping effort, and individual leeches / leech samples.

| | | | | | |
|---|---|---|---|---|---|
| *Crocidura sokolovi* | *Hystrix brachyura* | *Melogale moschata* | *Paguma larvata* | *Rattus losea* | *Viverra zibetha* |
| *Cuon alpinus* | *Leopoldamys edwardsi* | *Melogale personata* | *Panthera pardus* | *Rattus nitidus* | *Viverricula indica* |
| *Dendrogale murina* | *Leopoldamys sabanu* | *Muntiacus rooseveltorum* | *Panthera tigris,* | *Rattus tanezumi* | |
| *Dremomys rufigenis* | *Lepus peguensis* | *Muntiacus truongonensis* | *Paradoxuru hermaphroditus* | *Ratufa bicolor* | |
| *Elephas maximus* | *Lophura diardi* | *Muntiacus vaginalis* | *Pardofelis marmorata* | *Rheinardia ocellata* | |
| *Felis catus* | *Lophura edwardsi* | *Muntiacus vuquangensis* | *Pavo muticus* | *Rhinoceros sondaicus* | |
| *Galeopterus variegatus* | *Lophura nycthemera* | *Mus caroli* | *Petaurista elegans* | *Rhizomys pruinosus* | |
| *Gallus gallus* | *Lutra lutra* | *Mus cookii* | *Petaurista philippensis* | *Rhizomys sumatrensis* | |
| *Helarctos malayanus* | *Lutra sumatrana* | *Mus musculus* | *Pitta moluccensis* | *Rusa unicolor* | |
| *Herpestes javanicus* | *Lutrogale perspicillata* | *Mus pahari* | *Pitta nympha* | *Suncus murinus* | |
| *Herpestes urva* | *Macaca arctoides* | *Mustela kathiah* | *Polyplectron bicalcaratum* | *Sus scrofa* | |
| *Hydrornis cyaneus* | *Macaca assamensis* | *Mustela strigidorsa* | *Prionailurus bengalensis* | *Tamiops maritimus* | |
| *Hydrornis elliotii* | *Macaca fascicularis* | *Neofelis nebulosa* | *Prionodon pardicolor* | *Tamiops rodolphii* | |
| *Hydrornis oatesi* | *Macaca leonina* | *Nesolagus timminsi* | *Pseudoryx nghetinhensis* | *Trachypithecus laotum* | |
| *Hydrornis phayrei* | *Macaca mulatta* | *Niviventer fulvescens* | *Pygathrix cinerea* | *Trachypithecus phayrei* | |
| *Hydrornis soror* | *Manis javanica* | *Niviventer langbianis* | *Pygathrix nemaeus* | *Tragulus kanchil* | |
| *Hylomys suillus* | *Manis pentadactyla* | *Niviventer tenaster* | *Pygathrix nigripes* | *Tragulus versicolor* | |
| *Hylopetes alboniger* | *Martes flavigula* | *Nomascus annamensis* | *Rattus andamanensis* | *Tupaia belangeri* | |
| *Hylopetes phayrei* | *Maxomys moi* | *Nomascus gabriellae* | *Rattus argentiventer* | *Ursus thibetanus* | |
| *Hylopetes spadiceus* | *Maxomys surifer* | *Nycticebus bengalensis* | *Rattus exulans* | *Viverra megaspila* | |

**Table S2:** Species list to assign *a priori* weighting for PROTAX taxonomic assignment.

| | Species | | Detection method | | Conservation priority | |
|---|---|---|---|---|---|---|
| **No.** | **Common name** | **Scientific name** | **Camera traps** | **Leeches** | **IUCN Red List** | **Endemic** |
| 1 | Annamite striped rabbit | *Nesolagus timminsi* | X | X | EN | X |
| 2 | Asiatic black bear | *Ursus thibetanus* | X | X | VU | |
| 3 | Asiatic brush-tailed porcupine | *Atherurus macrourus* | X | X | LC | |
| 4 | Common palm civet | *Paradoxurus hermaphroditus* | X | X | LC | |
| 5 | Crab-eating mongoose | *Herpestes urva* | X | X | LC | |
| 6 | Dark muntjac complex | *Muntiacus rooseveltorum / truongsonensis* | X | X | DD | X |
| 7 | Eurasian wild pig | *Sus scrofa* | X | X | LC | |
| 8 | Ferret badger | *Melogale* spp. | X | X | LC | |
| 9 | Leopard cat | *Prionailurus bengalensis* | X | X | LC | |
| 10 | Malayan porcupine | *Hystrix brachyura* | X | X | LC | |
| 11 | Marbled cat | *Pardofelis marmorata* | | X | NT | |
| 12 | Masked palm civet | *Paguma larvata* | X | X | LC | |
| 13 | Northern treeshrew | *Tupaia belangeri* | X | X | LC | |
| 14 | Owston's civet | *Chrotogale owstoni* | X | X | EN | X |
| 15 | Pangolin | *Manis* spp. | X | | CR | |
| 16 | Pig-tailed macaque | *Macaca leonina* | X | | VU | |
| 17 | Red muntjac | *Muntiacus vaginalis* | X | X | LC | |
| 18 | Rhesus macaque | *Macaca mulatta* | X | X | LC | |
| 19 | Serow | *Capricornis milneedwardsii* | X | X | NT | |
| 20 | Spotted linsang | *Prionodon pardicolor* | X | | LC | |
| 21 | Stump-tailed macaque | *Macaca arctoides* | X | X | VU | |
| 22 | Yellow-bellied weasel | *Mustela kathiah* | X | | LC | |
| 23 | Yellow-throated marten | *Martes flavigula* | X | X | LC | |

**Table S3:** Final mammal species list for the multi-species occupancy model.

| | Mean | SD | Naive SE | Time-series SE | 2.50% | 25% | 50% | 75% | 97.50% | Point est. | Upper C.I. | BP value |
|---|---|---|---|---|---|---|---|---|---|---|---|---|
| R3 | 1265.4075 | 20.9444 | 0.0270 | 0.0546 | 1225.2244 | 1251.1491 | 1265.0806 | 1279.3409 | 1307.3547 | 1.0000 | 1.0000 | 0.5758 |
| alpha[1] | -1.4025 | 0.2511 | 0.0003 | 0.0006 | -1.9062 | -1.5685 | -1.3990 | -1.2317 | -0.9210 | 1.0000 | 1.0000 | 0.5758 |
| alpha[2] | -3.2270 | 1.0328 | 0.0013 | 0.0091 | -5.2408 | -3.8994 | -3.2446 | -2.5674 | -1.1273 | 1.0008 | 1.0028 | 0.5758 |
| alpha[3] | -0.7262 | 0.2267 | 0.0003 | 0.0005 | -1.1703 | -0.8785 | -0.7265 | -0.5746 | -0.2793 | 1.0000 | 1.0001 | 0.5758 |
| alpha[4] | -1.0056 | 0.2957 | 0.0004 | 0.0009 | -1.5719 | -1.2036 | -1.0111 | -0.8140 | -0.4071 | 1.0001 | 1.0001 | 0.5758 |
| alpha[5] | -0.3309 | 0.2712 | 0.0004 | 0.0009 | -0.8249 | -0.5150 | -0.3445 | -0.1631 | 0.2431 | 1.0000 | 1.0001 | 0.5758 |
| alpha[6] | -1.0463 | 0.4617 | 0.0006 | 0.0023 | -1.8980 | -1.3575 | -1.0669 | -0.7600 | -0.0748 | 1.0001 | 1.0001 | 0.5758 |
| alpha[7] | 0.4995 | 0.2608 | 0.0003 | 0.0008 | 0.0262 | 0.3204 | 0.4864 | 0.6627 | 1.0522 | 1.0001 | 1.0001 | 0.5758 |
| alpha[8] | 0.6396 | 0.2637 | 0.0003 | 0.0008 | 0.1611 | 0.4579 | 0.6258 | 0.8057 | 1.1966 | 1.0000 | 1.0001 | 0.5758 |
| alpha[9] | -0.8819 | 0.4508 | 0.0006 | 0.0025 | -1.6730 | -1.1903 | -0.9155 | -0.6115 | 0.1046 | 1.0000 | 1.0000 | 0.5758 |
| alpha[10] | -1.5738 | 0.5820 | 0.0008 | 0.0039 | -2.6103 | -1.9670 | -1.6112 | -1.2244 | -0.3158 | 1.0003 | 1.0009 | 0.5758 |
| alpha[11] | -3.2641 | 1.1046 | 0.0014 | 0.0108 | -5.3731 | -3.9893 | -3.3012 | -2.5725 | -0.9676 | 1.0009 | 1.0029 | 0.5758 |
| alpha[12] | -0.3937 | 0.6070 | 0.0008 | 0.0047 | -1.3741 | -0.8086 | -0.4647 | -0.0604 | 0.9958 | 1.0003 | 1.0008 | 0.5758 |
| alpha[13] | -1.3582 | 0.5858 | 0.0008 | 0.0045 | -2.3302 | -1.7530 | -1.4183 | -1.0353 | -0.0266 | 1.0018 | 1.0037 | 0.5758 |
| alpha[14] | -3.6186 | 1.1169 | 0.0014 | 0.0099 | -5.8014 | -4.3408 | -3.6383 | -2.9155 | -1.3301 | 1.0006 | 1.0019 | 0.5758 |
| alpha[15] | -2.6704 | 0.8642 | 0.0011 | 0.0075 | -4.3031 | -3.2425 | -2.7039 | -2.1277 | -0.8743 | 1.0009 | 1.0029 | 0.5758 |
| alpha[16] | -0.8966 | 0.2838 | 0.0004 | 0.0009 | -1.4423 | -1.0868 | -0.9009 | -0.7118 | -0.3244 | 1.0001 | 1.0003 | 0.5758 |
| alpha[17] | -1.3567 | 0.2815 | 0.0004 | 0.0008 | -1.9133 | -1.5438 | -1.3546 | -1.1683 | -0.8071 | 1.0000 | 1.0001 | 0.5758 |
| alpha[18] | -2.7496 | 0.7939 | 0.0010 | 0.0062 | -4.2415 | -3.2773 | -2.7808 | -2.2565 | -1.0811 | 1.0003 | 1.0010 | 0.5758 |
| alpha[19] | -0.9661 | 0.2911 | 0.0004 | 0.0009 | -1.5198 | -1.1623 | -0.9725 | -0.7782 | -0.3736 | 1.0001 | 1.0002 | 0.5758 |
| alpha[20] | -0.6948 | 0.6110 | 0.0008 | 0.0048 | -1.6920 | -1.1143 | -0.7626 | -0.3551 | 0.7062 | 1.0013 | 1.0025 | 0.5758 |
| alpha[21] | 0.6542 | 0.3012 | 0.0004 | 0.0010 | 0.1087 | 0.4461 | 0.6387 | 0.8452 | 1.2892 | 1.0002 | 1.0005 | 0.5758 |
| alpha[22] | -2.8491 | 0.9259 | 0.0012 | 0.0085 | -4.5796 | -3.4643 | -2.8891 | -2.2749 | -0.8921 | 1.0011 | 1.0032 | 0.5758 |
| alpha[23] | -0.9102 | 0.5688 | 0.0007 | 0.0042 | -1.8551 | -1.3000 | -0.9688 | -0.5884 | 0.3789 | 1.0001 | 1.0001 | 0.5758 |
| b.CT | 0.6531 | 0.0778 | 0.0001 | 0.0006 | 0.5031 | 0.6001 | 0.6522 | 0.7052 | 0.8081 | 1.0001 | 1.0002 | 0.5758 |

| | | | | | | | | | | | |
|---|---|---|---|---|---|---|---|---|---|---|---|
| b.eff | 0.0228 | 11.548 0 | 0.0149 | 0.0149 | -18.9957 | -9.9761 | 0.0454 | 10.0338 | 19.0059 | 1.0000 | 1.0000 | 0.5758 |
| b.leech | 0.2097 | 0.0325 | 0.0000 | 0.0001 | 0.1477 | 0.1876 | 0.2092 | 0.2313 | 0.2751 | 1.0000 | 1.0000 | 0.5758 |
| beta1[1] | 0.2455 | 0.2867 | 0.0004 | 0.0008 | -0.3113 | 0.0520 | 0.2437 | 0.4369 | 0.8134 | 1.0000 | 1.0001 | 0.5758 |
| beta1[2] | -0.1246 | 0.6365 | 0.0008 | 0.0030 | -1.3831 | -0.5338 | -0.1228 | 0.2827 | 1.1337 | 1.0001 | 1.0003 | 0.5758 |
| beta1[3] | 0.1095 | 0.2656 | 0.0003 | 0.0007 | -0.4098 | -0.0688 | 0.1079 | 0.2860 | 0.6355 | 1.0000 | 1.0001 | 0.5758 |
| beta1[4] | 0.3403 | 0.2976 | 0.0004 | 0.0008 | -0.2357 | 0.1401 | 0.3373 | 0.5367 | 0.9356 | 1.0000 | 1.0000 | 0.5758 |
| beta1[5] | 0.4055 | 0.3069 | 0.0004 | 0.0011 | -0.1645 | 0.1975 | 0.3937 | 0.6002 | 1.0438 | 1.0000 | 1.0001 | 0.5758 |
| beta1[6] | 0.9880 | 0.5019 | 0.0006 | 0.0023 | 0.1026 | 0.6440 | 0.9513 | 1.2921 | 2.0830 | 1.0000 | 1.0001 | 0.5758 |
| beta1[7] | 0.0582 | 0.2818 | 0.0004 | 0.0008 | -0.4970 | -0.1293 | 0.0586 | 0.2449 | 0.6142 | 1.0000 | 1.0001 | 0.5758 |
| beta1[8] | 0.7277 | 0.2928 | 0.0004 | 0.0009 | 0.1748 | 0.5285 | 0.7199 | 0.9177 | 1.3270 | 1.0000 | 1.0001 | 0.5758 |
| beta1[9] | -1.0749 | 0.4433 | 0.0006 | 0.0021 | -2.0342 | -1.3451 | -1.0443 | -0.7705 | -0.2914 | 1.0002 | 1.0005 | 0.5758 |
| beta1[10] | -0.9108 | 0.4778 | 0.0006 | 0.0021 | -1.9278 | -1.2075 | -0.8842 | -0.5837 | -0.0494 | 1.0001 | 1.0001 | 0.5758 |
| beta1[11] | -0.0374 | 0.6497 | 0.0008 | 0.0028 | -1.3506 | -0.4504 | -0.0271 | 0.3867 | 1.2134 | 1.0001 | 1.0004 | 0.5758 |
| beta1[12] | 0.4810 | 0.5061 | 0.0007 | 0.0027 | -0.4314 | 0.1446 | 0.4491 | 0.7809 | 1.5810 | 1.0000 | 1.0000 | 0.5758 |
| beta1[13] | -0.0229 | 0.4226 | 0.0005 | 0.0018 | -0.8574 | -0.2970 | -0.0221 | 0.2512 | 0.8111 | 1.0001 | 1.0001 | 0.5758 |
| beta1[14] | 0.5330 | 0.6278 | 0.0008 | 0.0025 | -0.6585 | 0.1166 | 0.5145 | 0.9300 | 1.8260 | 1.0001 | 1.0001 | 0.5758 |
| beta1[15] | 0.0114 | 0.5707 | 0.0007 | 0.0026 | -1.0888 | -0.3617 | 0.0029 | 0.3723 | 1.1714 | 1.0002 | 1.0006 | 0.5758 |
| beta1[16] | 0.4602 | 0.3385 | 0.0004 | 0.0012 | -0.1787 | 0.2297 | 0.4511 | 0.6813 | 1.1489 | 1.0002 | 1.0009 | 0.5758 |
| beta1[17] | -0.6214 | 0.3178 | 0.0004 | 0.0010 | -1.2669 | -0.8301 | -0.6135 | -0.4051 | -0.0188 | 1.0001 | 1.0003 | 0.5758 |
| beta1[18] | -0.9940 | 0.6063 | 0.0008 | 0.0029 | -2.3050 | -1.3638 | -0.9538 | -0.5771 | 0.0792 | 1.0000 | 1.0001 | 0.5758 |
| beta1[19] | 0.5356 | 0.3131 | 0.0004 | 0.0010 | -0.0525 | 0.3226 | 0.5254 | 0.7376 | 1.1787 | 1.0000 | 1.0001 | 0.5758 |
| beta1[20] | 1.0135 | 0.4708 | 0.0006 | 0.0023 | 0.1769 | 0.6948 | 0.9808 | 1.2967 | 2.0361 | 1.0001 | 1.0002 | 0.5758 |
| beta1[21] | 1.8522 | 0.4277 | 0.0006 | 0.0018 | 1.0806 | 1.5548 | 1.8280 | 2.1237 | 2.7579 | 1.0001 | 1.0005 | 0.5758 |
| beta1[22] | 0.5018 | 0.6124 | 0.0008 | 0.0029 | -0.6225 | 0.0969 | 0.4716 | 0.8703 | 1.8076 | 1.0001 | 1.0001 | 0.5758 |
| beta1[23] | 0.3026 | 0.4108 | 0.0005 | 0.0018 | -0.4827 | 0.0299 | 0.2939 | 0.5659 | 1.1399 | 1.0001 | 1.0002 | 0.5758 |
| beta2[1] | -0.2611 | 0.2355 | 0.0003 | 0.0006 | -0.7275 | -0.4160 | -0.2613 | -0.1061 | 0.2040 | 1.0000 | 1.0000 | 0.5758 |

| | | | | | | | | | | | | |
|---|---|---|---|---|---|---|---|---|---|---|---|---|
| beta2[2] | -0.4176 | 0.3463 | 0.0004 | 0.0011 | -1.1548 | -0.6253 | -0.4021 | -0.1933 | 0.2324 | 1.0000 | 1.0000 | 0.5758 |
| beta2[3] | -0.4563 | 0.2097 | 0.0003 | 0.0006 | -0.8835 | -0.5923 | -0.4508 | -0.3146 | -0.0589 | 1.0000 | 1.0001 | 0.5758 |
| beta2[4] | -0.2145 | 0.2336 | 0.0003 | 0.0007 | -0.6719 | -0.3682 | -0.2163 | -0.0625 | 0.2512 | 1.0000 | 1.0000 | 0.5758 |
| beta2[5] | -0.1883 | 0.2173 | 0.0003 | 0.0006 | -0.6112 | -0.3328 | -0.1900 | -0.0456 | 0.2453 | 1.0001 | 1.0002 | 0.5758 |
| beta2[6] | -0.5818 | 0.3109 | 0.0004 | 0.0011 | -1.2573 | -0.7697 | -0.5593 | -0.3703 | -0.0294 | 1.0001 | 1.0002 | 0.5758 |
| beta2[7] | -0.6063 | 0.2426 | 0.0003 | 0.0008 | -1.1239 | -0.7569 | -0.5915 | -0.4400 | -0.1696 | 1.0001 | 1.0003 | 0.5758 |
| beta2[8] | -0.4046 | 0.2164 | 0.0003 | 0.0006 | -0.8464 | -0.5448 | -0.3994 | -0.2591 | 0.0075 | 1.0000 | 1.0001 | 0.5758 |
| beta2[9] | -0.2802 | 0.2680 | 0.0003 | 0.0009 | -0.8168 | -0.4514 | -0.2789 | -0.1075 | 0.2495 | 1.0000 | 1.0001 | 0.5758 |
| beta2[10] | -0.5152 | 0.2982 | 0.0004 | 0.0011 | -1.1497 | -0.6983 | -0.4995 | -0.3156 | 0.0307 | 1.0001 | 1.0005 | 0.5758 |
| beta2[11] | -0.3097 | 0.3400 | 0.0004 | 0.0010 | -0.9978 | -0.5214 | -0.3082 | -0.0961 | 0.3638 | 1.0000 | 1.0001 | 0.5758 |
| beta2[12] | -0.2659 | 0.2824 | 0.0004 | 0.0009 | -0.8278 | -0.4477 | -0.2657 | -0.0837 | 0.2943 | 1.0001 | 1.0002 | 0.5758 |
| beta2[13] | -0.3482 | 0.2869 | 0.0004 | 0.0009 | -0.9376 | -0.5278 | -0.3423 | -0.1612 | 0.2035 | 1.0000 | 1.0000 | 0.5758 |
| beta2[14] | -0.3541 | 0.3424 | 0.0004 | 0.0009 | -1.0614 | -0.5638 | -0.3463 | -0.1359 | 0.3108 | 1.0000 | 1.0000 | 0.5758 |
| beta2[15] | -0.4720 | 0.3444 | 0.0004 | 0.0011 | -1.2144 | -0.6779 | -0.4525 | -0.2456 | 0.1600 | 1.0001 | 1.0003 | 0.5758 |
| beta2[16] | 0.0736 | 0.2440 | 0.0003 | 0.0008 | -0.3868 | -0.0925 | 0.0663 | 0.2319 | 0.5729 | 1.0000 | 1.0001 | 0.5758 |
| beta2[17] | -0.5740 | 0.2585 | 0.0003 | 0.0008 | -1.1204 | -0.7375 | -0.5601 | -0.3967 | -0.1030 | 1.0001 | 1.0002 | 0.5758 |
| beta2[18] | -0.4065 | 0.3227 | 0.0004 | 0.0010 | -1.0864 | -0.6037 | -0.3944 | -0.1961 | 0.2017 | 1.0000 | 1.0000 | 0.5758 |
| beta2[19] | -0.0645 | 0.2449 | 0.0003 | 0.0007 | -0.5268 | -0.2302 | -0.0719 | 0.0936 | 0.4381 | 1.0000 | 1.0001 | 0.5758 |
| beta2[20] | -0.1018 | 0.2762 | 0.0004 | 0.0010 | -0.6226 | -0.2859 | -0.1112 | 0.0722 | 0.4712 | 1.0000 | 1.0000 | 0.5758 |
| beta2[21] | -0.2640 | 0.2330 | 0.0003 | 0.0006 | -0.7196 | -0.4177 | -0.2656 | -0.1118 | 0.2021 | 1.0000 | 1.0001 | 0.5758 |
| beta2[22] | -0.3240 | 0.3335 | 0.0004 | 0.0010 | -1.0063 | -0.5304 | -0.3203 | -0.1125 | 0.3313 | 1.0001 | 1.0004 | 0.5758 |
| beta2[23] | -0.2679 | 0.2673 | 0.0003 | 0.0009 | -0.7970 | -0.4414 | -0.2682 | -0.0949 | 0.2627 | 1.0001 | 1.0002 | 0.5758 |
| beta3[1] | 0.3237 | 0.2967 | 0.0004 | 0.0008 | -0.2378 | 0.1212 | 0.3165 | 0.5187 | 0.9269 | 1.0000 | 1.0000 | 0.5758 |
| beta3[2] | 0.4947 | 0.6071 | 0.0008 | 0.0028 | -0.5985 | 0.0787 | 0.4584 | 0.8714 | 1.7936 | 1.0001 | 1.0004 | 0.5758 |
| beta3[3] | -0.8948 | 0.2820 | 0.0004 | 0.0009 | -1.4736 | -1.0777 | -0.8860 | -0.7018 | -0.3664 | 1.0000 | 1.0002 | 0.5758 |
| beta3[4] | -1.0320 | 0.3286 | 0.0004 | 0.0011 | -1.7188 | -1.2406 | -1.0186 | -0.8063 | -0.4276 | 1.0000 | 1.0002 | 0.5758 |
| beta3[5] | -0.1311 | 0.3042 | 0.0004 | 0.0010 | -0.7445 | -0.3289 | -0.1267 | 0.0719 | 0.4565 | 1.0000 | 1.0001 | 0.5758 |

| | | | | | | | | | | | |
|---|---|---|---|---|---|---|---|---|---|---|---|
| beta3[6] | -0.1154 | 0.4319 | 0.0006 | 0.0018 | -0.9917 | -0.3939 | -0.1067 | 0.1731 | 0.7151 | 1.0000 | 1.0001 | 0.5758 |
| beta3[7] | -0.2472 | 0.3044 | 0.0004 | 0.0010 | -0.8615 | -0.4451 | -0.2433 | -0.0432 | 0.3394 | 1.0000 | 1.0000 | 0.5758 |
| beta3[8] | -0.0414 | 0.2822 | 0.0004 | 0.0008 | -0.6000 | -0.2278 | -0.0396 | 0.1468 | 0.5090 | 1.0000 | 1.0001 | 0.5758 |
| beta3[9] | 0.0384 | 0.4077 | 0.0005 | 0.0018 | -0.7555 | -0.2251 | 0.0350 | 0.2996 | 0.8518 | 1.0000 | 1.0001 | 0.5758 |
| beta3[10] | -0.0867 | 0.4337 | 0.0006 | 0.0018 | -0.9344 | -0.3697 | -0.0895 | 0.1940 | 0.7765 | 1.0001 | 1.0001 | 0.5758 |
| beta3[11] | 0.0063 | 0.6020 | 0.0008 | 0.0024 | -1.1499 | -0.3889 | -0.0068 | 0.3870 | 1.2352 | 1.0000 | 1.0001 | 0.5758 |
| beta3[12] | 0.4815 | 0.4405 | 0.0006 | 0.0020 | -0.3616 | 0.1899 | 0.4723 | 0.7631 | 1.3779 | 1.0000 | 1.0001 | 0.5758 |
| beta3[13] | 0.1765 | 0.4224 | 0.0005 | 0.0018 | -0.6306 | -0.1033 | 0.1667 | 0.4461 | 1.0346 | 1.0000 | 1.0001 | 0.5758 |
| beta3[14] | 0.0650 | 0.5921 | 0.0008 | 0.0022 | -1.0539 | -0.3281 | 0.0448 | 0.4384 | 1.2921 | 1.0002 | 1.0003 | 0.5758 |
| beta3[15] | 0.7461 | 0.5999 | 0.0008 | 0.0030 | -0.3152 | 0.3319 | 0.7056 | 1.1155 | 2.0406 | 1.0002 | 1.0004 | 0.5758 |
| beta3[16] | -1.2966 | 0.3906 | 0.0005 | 0.0016 | -2.1393 | -1.5384 | -1.2695 | -1.0254 | -0.6069 | 1.0003 | 1.0009 | 0.5758 |
| beta3[17] | 0.4535 | 0.3353 | 0.0004 | 0.0011 | -0.1752 | 0.2233 | 0.4435 | 0.6725 | 1.1400 | 1.0001 | 1.0003 | 0.5758 |
| beta3[18] | 0.3319 | 0.5548 | 0.0007 | 0.0025 | -0.7022 | -0.0412 | 0.3102 | 0.6815 | 1.4907 | 1.0001 | 1.0003 | 0.5758 |
| beta3[19] | -0.1549 | 0.3160 | 0.0004 | 0.0010 | -0.7890 | -0.3624 | -0.1514 | 0.0581 | 0.4557 | 1.0000 | 1.0000 | 0.5758 |
| beta3[20] | -0.3799 | 0.3782 | 0.0005 | 0.0015 | -1.1403 | -0.6245 | -0.3760 | -0.1295 | 0.3534 | 1.0001 | 1.0001 | 0.5758 |
| beta3[21] | -1.2051 | 0.3748 | 0.0005 | 0.0016 | -1.9895 | -1.4461 | -1.1876 | -0.9458 | -0.5187 | 1.0001 | 1.0003 | 0.5758 |
| beta3[22] | 0.3998 | 0.5781 | 0.0007 | 0.0025 | -0.6472 | 0.0069 | 0.3672 | 0.7572 | 1.6299 | 1.0001 | 1.0003 | 0.5758 |
| beta3[23] | -0.4450 | 0.4178 | 0.0005 | 0.0019 | -1.3167 | -0.7066 | -0.4287 | -0.1650 | 0.3347 | 1.0000 | 1.0002 | 0.5758 |
| beta4[1] | -0.1771 | 0.2250 | 0.0003 | 0.0006 | -0.6294 | -0.3256 | -0.1731 | -0.0250 | 0.2557 | 1.0000 | 1.0001 | 0.5758 |
| beta4[2] | 0.3350 | 0.3581 | 0.0005 | 0.0013 | -0.3180 | 0.0934 | 0.3153 | 0.5558 | 1.0983 | 1.0000 | 1.0001 | 0.5758 |
| beta4[3] | 0.0514 | 0.2213 | 0.0003 | 0.0005 | -0.3856 | -0.0956 | 0.0518 | 0.1990 | 0.4863 | 1.0000 | 1.0000 | 0.5758 |
| beta4[4] | -0.2375 | 0.2672 | 0.0003 | 0.0007 | -0.7912 | -0.4099 | -0.2273 | -0.0547 | 0.2582 | 1.0000 | 1.0000 | 0.5758 |
| beta4[5] | 0.0286 | 0.2330 | 0.0003 | 0.0006 | -0.4225 | -0.1259 | 0.0259 | 0.1796 | 0.4974 | 1.0000 | 1.0001 | 0.5758 |
| beta4[6] | 0.3006 | 0.2903 | 0.0004 | 0.0009 | -0.2404 | 0.1049 | 0.2890 | 0.4842 | 0.9051 | 1.0000 | 1.0002 | 0.5758 |
| beta4[7] | 0.5165 | 0.2685 | 0.0003 | 0.0009 | 0.0375 | 0.3314 | 0.4995 | 0.6837 | 1.0912 | 1.0000 | 1.0001 | 0.5758 |
| beta4[8] | -0.4865 | 0.2525 | 0.0003 | 0.0008 | -1.0068 | -0.6496 | -0.4781 | -0.3141 | -0.0140 | 1.0000 | 1.0001 | 0.5758 |
| beta4[9] | 0.3114 | 0.2973 | 0.0004 | 0.0010 | -0.2349 | 0.1103 | 0.2972 | 0.4967 | 0.9383 | 1.0000 | 1.0000 | 0.5758 |

| | | | | | | | | | | | |
|---|---|---|---|---|---|---|---|---|---|---|---|
| beta4[10] | 0.0187 | 0.3117 | 0.0004 | 0.0010 | -0.6114 | -0.1805 | 0.0226 | 0.2218 | 0.6264 | 1.0000 | 1.0000 | 0.5758 |
| beta4[11] | -0.1587 | 0.3850 | 0.0005 | 0.0012 | -0.9794 | -0.3922 | -0.1397 | 0.0957 | 0.5510 | 1.0000 | 1.0001 | 0.5758 |
| beta4[12] | -0.3250 | 0.3139 | 0.0004 | 0.0012 | -0.9695 | -0.5242 | -0.3154 | -0.1168 | 0.2722 | 1.0001 | 1.0002 | 0.5758 |
| beta4[13] | -0.0856 | 0.3046 | 0.0004 | 0.0011 | -0.6927 | -0.2813 | -0.0835 | 0.1110 | 0.5146 | 1.0001 | 1.0001 | 0.5758 |
| beta4[14] | -0.0838 | 0.3742 | 0.0005 | 0.0012 | -0.8551 | -0.3169 | -0.0747 | 0.1584 | 0.6367 | 1.0000 | 1.0000 | 0.5758 |
| beta4[15] | 0.0198 | 0.3544 | 0.0005 | 0.0013 | -0.6709 | -0.2083 | 0.0149 | 0.2408 | 0.7443 | 1.0000 | 1.0001 | 0.5758 |
| beta4[16] | 0.0072 | 0.2513 | 0.0003 | 0.0006 | -0.4942 | -0.1588 | 0.0095 | 0.1749 | 0.4958 | 1.0000 | 1.0000 | 0.5758 |
| beta4[17] | 0.2482 | 0.2388 | 0.0003 | 0.0006 | -0.2045 | 0.0862 | 0.2420 | 0.4025 | 0.7360 | 1.0000 | 1.0000 | 0.5758 |
| beta4[18] | 0.0161 | 0.3449 | 0.0004 | 0.0010 | -0.6809 | -0.2034 | 0.0196 | 0.2391 | 0.6932 | 1.0000 | 1.0001 | 0.5758 |
| beta4[19] | 0.4080 | 0.2436 | 0.0003 | 0.0007 | -0.0429 | 0.2409 | 0.3984 | 0.5650 | 0.9139 | 1.0000 | 1.0000 | 0.5758 |
| beta4[20] | -0.2126 | 0.3323 | 0.0004 | 0.0013 | -0.8897 | -0.4231 | -0.2057 | 0.0041 | 0.4292 | 1.0000 | 1.0001 | 0.5758 |
| beta4[21] | 0.2508 | 0.2635 | 0.0003 | 0.0007 | -0.2467 | 0.0732 | 0.2429 | 0.4201 | 0.7932 | 1.0000 | 1.0000 | 0.5758 |
| beta4[22] | -0.0848 | 0.3611 | 0.0005 | 0.0013 | -0.8170 | -0.3124 | -0.0802 | 0.1461 | 0.6247 | 1.0000 | 1.0000 | 0.5758 |
| beta4[23] | 0.1152 | 0.2966 | 0.0004 | 0.0010 | -0.4543 | -0.0794 | 0.1095 | 0.3027 | 0.7209 | 1.0000 | 1.0000 | 0.5758 |
| mu.a | -1.3104 | 0.3674 | 0.0005 | 0.0026 | -2.0724 | -1.5455 | -1.2966 | -1.0600 | -0.6291 | 1.0005 | 1.0014 | 0.5758 |
| mu.b1 | 0.2076 | 0.2118 | 0.0003 | 0.0007 | -0.2139 | 0.0713 | 0.2080 | 0.3452 | 0.6259 | 1.0000 | 1.0000 | 0.5758 |
| mu.b2 | -0.3306 | 0.1187 | 0.0002 | 0.0005 | -0.5715 | -0.4071 | -0.3284 | -0.2515 | -0.1029 | 1.0001 | 1.0003 | 0.5758 |
| mu.b3 | -0.1093 | 0.1955 | 0.0003 | 0.0007 | -0.4793 | -0.2398 | -0.1151 | 0.0145 | 0.2939 | 1.0000 | 1.0000 | 0.5758 |
| mu.b4 | 0.0338 | 0.1235 | 0.0002 | 0.0004 | -0.2112 | -0.0467 | 0.0342 | 0.1147 | 0.2770 | 1.0000 | 1.0000 | 0.5758 |
| mu.bp[1] | -3.0597 | 0.2892 | 0.0004 | 0.0023 | -3.6962 | -3.2333 | -3.0359 | -2.8602 | -2.5581 | 1.0007 | 1.0018 | 0.5758 |
| mu.bp[2] | -3.2938 | 0.4288 | 0.0006 | 0.0023 | -4.2325 | -3.5524 | -3.2613 | -2.9994 | -2.5447 | 1.0003 | 1.0010 | 0.5758 |
| mu.bp[3] | -2.5342 | 0.4300 | 0.0006 | 0.0034 | -3.5389 | -2.7590 | -2.4803 | -2.2445 | -1.8521 | 1.0008 | 1.0018 | 0.5758 |
| mu.p[1,1] | -1.9146 | 0.2106 | 0.0003 | 0.0009 | -2.3375 | -2.0541 | -1.9111 | -1.7714 | -1.5120 | 1.0000 | 1.0001 | 0.5758 |
| mu.p[2,1] | -4.1779 | 0.9066 | 0.0012 | 0.0079 | -6.1681 | -4.7393 | -4.0957 | -3.5325 | -2.6325 | 1.0010 | 1.0030 | 0.5758 |
| mu.p[3,1] | -1.9623 | 0.1816 | 0.0002 | 0.0009 | -2.3233 | -2.0836 | -1.9605 | -1.8386 | -1.6121 | 1.0000 | 1.0002 | 0.5758 |
| mu.p[4,1] | -2.4240 | 0.2333 | 0.0003 | 0.0010 | -2.8922 | -2.5791 | -2.4199 | -2.2644 | -1.9784 | 1.0001 | 1.0002 | 0.5758 |
| mu.p[5,1] | -2.3341 | 0.2019 | 0.0003 | 0.0010 | -2.7407 | -2.4679 | -2.3304 | -2.1964 | -1.9488 | 1.0000 | 1.0001 | 0.5758 |

| | | | | | | | | | | | | |
|---|---|---|---|---|---|---|---|---|---|---|---|---|
| mu.p[6,1] | -3.5670 | 0.3170 | 0.0004 | 0.0014 | -4.2083 | -3.7765 | -3.5594 | -3.3492 | -2.9664 | 1.0000 | 1.0000 | 0.5758 |
| mu.p[7,1] | -2.1327 | 0.1574 | 0.0002 | 0.0009 | -2.4453 | -2.2382 | -2.1312 | -2.0259 | -1.8285 | 1.0001 | 1.0003 | 0.5758 |
| mu.p[8,1] | -2.0544 | 0.1475 | 0.0002 | 0.0008 | -2.3465 | -2.1533 | -2.0531 | -1.9542 | -1.7690 | 1.0000 | 1.0001 | 0.5758 |
| mu.p[9,1] | -3.0855 | 0.3153 | 0.0004 | 0.0016 | -3.7189 | -3.2966 | -3.0793 | -2.8687 | -2.4857 | 1.0000 | 1.0001 | 0.5758 |
| mu.p[10,1] | -3.5103 | 0.4564 | 0.0006 | 0.0029 | -4.4430 | -3.8121 | -3.4955 | -3.1929 | -2.6613 | 1.0002 | 1.0005 | 0.5758 |
| mu.p[11,1] | -4.3996 | 1.0738 | 0.0014 | 0.0100 | -6.7915 | -5.0521 | -4.2823 | -3.6314 | -2.6227 | 1.0010 | 1.0030 | 0.5758 |
| mu.p[12,1] | -3.6050 | 0.3507 | 0.0005 | 0.0022 | -4.3010 | -3.8412 | -3.6016 | -3.3647 | -2.9309 | 1.0001 | 1.0005 | 0.5758 |
| mu.p[13,1] | -3.3443 | 0.4630 | 0.0006 | 0.0031 | -4.3145 | -3.6459 | -3.3213 | -3.0178 | -2.5037 | 1.0005 | 1.0015 | 0.5758 |
| mu.p[14,1] | -3.7605 | 1.0320 | 0.0013 | 0.0089 | -5.9910 | -4.4072 | -3.6752 | -3.0332 | -1.9560 | 1.0008 | 1.0026 | 0.5758 |
| mu.p[15,1] | -3.8589 | 0.7014 | 0.0009 | 0.0057 | -5.3367 | -4.3148 | -3.8157 | -3.3617 | -2.6101 | 1.0009 | 1.0028 | 0.5758 |
| mu.p[16,1] | -2.2717 | 0.2168 | 0.0003 | 0.0010 | -2.7069 | -2.4158 | -2.2682 | -2.1236 | -1.8578 | 1.0001 | 1.0003 | 0.5758 |
| mu.p[17,1] | -2.3077 | 0.2219 | 0.0003 | 0.0009 | -2.7527 | -2.4551 | -2.3038 | -2.1563 | -1.8839 | 1.0000 | 1.0002 | 0.5758 |
| mu.p[18,1] | -3.8961 | 0.7354 | 0.0009 | 0.0056 | -5.4643 | -4.3656 | -3.8439 | -3.3750 | -2.5966 | 1.0004 | 1.0014 | 0.5758 |
| mu.p[19,1] | -2.5947 | 0.2370 | 0.0003 | 0.0010 | -3.0734 | -2.7518 | -2.5894 | -2.4325 | -2.1447 | 1.0001 | 1.0002 | 0.5758 |
| mu.p[20,1] | -3.5254 | 0.3888 | 0.0005 | 0.0026 | -4.3054 | -3.7871 | -3.5177 | -3.2557 | -2.7884 | 1.0002 | 1.0007 | 0.5758 |
| mu.p[21,1] | -2.3196 | 0.1569 | 0.0002 | 0.0008 | -2.6315 | -2.4247 | -2.3180 | -2.2132 | -2.0163 | 1.0001 | 1.0003 | 0.5758 |
| mu.p[22,1] | -3.8338 | 0.8103 | 0.0010 | 0.0071 | -5.5427 | -4.3610 | -3.7757 | -3.2538 | -2.4075 | 1.0009 | 1.0031 | 0.5758 |
| mu.p[23,1] | -3.5178 | 0.3996 | 0.0005 | 0.0025 | -4.3320 | -3.7829 | -3.5048 | -3.2393 | -2.7728 | 1.0000 | 1.0001 | 0.5758 |
| mu.p[1,2] | -3.0987 | 0.6062 | 0.0008 | 0.0014 | -4.3863 | -3.4812 | -3.0642 | -2.6779 | -2.0083 | 1.0000 | 1.0001 | 0.5758 |
| mu.p[2,2] | -3.2514 | 1.0279 | 0.0013 | 0.0046 | -5.3994 | -3.8941 | -3.2124 | -2.5650 | -1.3258 | 1.0002 | 1.0007 | 0.5758 |
| mu.p[3,2] | -2.8821 | 0.5331 | 0.0007 | 0.0011 | -4.0087 | -3.2206 | -2.8543 | -2.5125 | -1.9150 | 1.0001 | 1.0002 | 0.5758 |
| mu.p[4,2] | -3.7682 | 0.8286 | 0.0011 | 0.0022 | -5.6125 | -4.2641 | -3.6945 | -3.1892 | -2.3557 | 1.0000 | 1.0001 | 0.5758 |
| mu.p[5,2] | -3.7277 | 0.5984 | 0.0008 | 0.0016 | -5.0099 | -4.1026 | -3.6899 | -3.3111 | -2.6625 | 1.0000 | 1.0001 | 0.5758 |
| mu.p[6,2] | -1.7422 | 0.4193 | 0.0005 | 0.0015 | -2.5763 | -2.0230 | -1.7384 | -1.4586 | -0.9310 | 1.0000 | 1.0000 | 0.5758 |
| mu.p[7,2] | -1.3614 | 0.2762 | 0.0004 | 0.0008 | -1.9153 | -1.5456 | -1.3567 | -1.1733 | -0.8316 | 1.0000 | 1.0001 | 0.5758 |
| mu.p[8,2] | -2.2898 | 0.3402 | 0.0004 | 0.0010 | -2.9845 | -2.5129 | -2.2806 | -2.0564 | -1.6491 | 1.0000 | 1.0000 | 0.5758 |
| mu.p[9,2] | -4.5000 | 1.1227 | 0.0014 | 0.0041 | -7.1088 | -5.1232 | -4.3578 | -3.7159 | -2.7189 | 1.0001 | 1.0002 | 0.5758 |

| | | | | | | | | | | | | |
|---|---|---|---|---|---|---|---|---|---|---|---|---|
| mu.p[10,2] | -3.5636 | 0.8729 | 0.0011 | 0.0028 | -5.4744 | -4.0959 | -3.4973 | -2.9587 | -2.0386 | 1.0001 | 1.0002 | 0.5758 |
| mu.p[11,2] | -2.7942 | 1.1353 | 0.0015 | 0.0059 | -5.0767 | -3.5184 | -2.7870 | -2.0628 | -0.5567 | 1.0002 | 1.0008 | 0.5758 |
| mu.p[12,2] | -3.0678 | 0.5438 | 0.0007 | 0.0021 | -4.1862 | -3.4219 | -3.0496 | -2.6945 | -2.0529 | 1.0001 | 1.0002 | 0.5758 |
| mu.p[13,2] | -4.4686 | 1.1594 | 0.0015 | 0.0048 | -7.1243 | -5.1205 | -4.3332 | -3.6620 | -2.5934 | 1.0002 | 1.0007 | 0.5758 |
| mu.p[14,2] | -3.9641 | 1.2764 | 0.0016 | 0.0057 | -6.8101 | -4.6998 | -3.8463 | -3.0951 | -1.7875 | 1.0002 | 1.0005 | 0.5758 |
| mu.p[15,2] | -4.2355 | 1.2445 | 0.0016 | 0.0057 | -7.0462 | -4.9512 | -4.1116 | -3.3792 | -2.1496 | 1.0003 | 1.0010 | 0.5758 |
| mu.p[16,2] | -4.6000 | 1.0862 | 0.0014 | 0.0037 | -7.1475 | -5.1893 | -4.4540 | -3.8411 | -2.9067 | 1.0001 | 1.0003 | 0.5758 |
| mu.p[17,2] | -2.4359 | 0.5121 | 0.0007 | 0.0011 | -3.5040 | -2.7665 | -2.4141 | -2.0817 | -1.4910 | 1.0000 | 1.0000 | 0.5758 |
| mu.p[18,2] | -2.1193 | 1.0022 | 0.0013 | 0.0044 | -4.0478 | -2.7886 | -2.1405 | -1.4657 | -0.0927 | 1.0001 | 1.0005 | 0.5758 |
| mu.p[19,2] | -2.0962 | 0.4479 | 0.0006 | 0.0012 | -3.0037 | -2.3915 | -2.0853 | -1.7898 | -1.2484 | 1.0000 | 1.0000 | 0.5758 |
| mu.p[20,2] | -4.8071 | 1.1091 | 0.0014 | 0.0046 | -7.3712 | -5.4234 | -4.6727 | -4.0336 | -3.0365 | 1.0001 | 1.0003 | 0.5758 |
| mu.p[21,2] | -2.8349 | 0.3991 | 0.0005 | 0.0010 | -3.6581 | -3.0945 | -2.8202 | -2.5598 | -2.0944 | 1.0000 | 1.0001 | 0.5758 |
| mu.p[22,2] | -4.2127 | 1.2506 | 0.0016 | 0.0061 | -7.0243 | -4.9375 | -4.0920 | -3.3514 | -2.1064 | 1.0003 | 1.0010 | 0.5758 |
| mu.p[23,2] | -3.9864 | 0.8618 | 0.0011 | 0.0032 | -5.8836 | -4.5068 | -3.9170 | -3.3859 | -2.5016 | 1.0000 | 1.0001 | 0.5758 |
| mu.p[1,3] | -2.3155 | 0.5711 | 0.0007 | 0.0017 | -3.5433 | -2.6557 | -2.2865 | -1.9419 | -1.2555 | 1.0000 | 1.0001 | 0.5758 |
| mu.p[2,3] | -2.6841 | 0.9208 | 0.0012 | 0.0050 | -4.9100 | -3.1038 | -2.5478 | -2.1101 | -1.2384 | 1.0006 | 1.0010 | 0.5758 |
| mu.p[3,3] | -2.8535 | 0.9097 | 0.0012 | 0.0055 | -5.1013 | -3.2588 | -2.6887 | -2.2622 | -1.5508 | 1.0005 | 1.0008 | 0.5758 |
| mu.p[4,3] | -2.6897 | 0.9212 | 0.0012 | 0.0049 | -4.9154 | -3.0988 | -2.5516 | -2.1168 | -1.2675 | 1.0005 | 1.0009 | 0.5758 |
| mu.p[5,3] | -2.3519 | 0.5688 | 0.0007 | 0.0017 | -3.5880 | -2.6869 | -2.3188 | -1.9771 | -1.3147 | 1.0000 | 1.0001 | 0.5758 |
| mu.p[6,3] | -2.8818 | 0.9202 | 0.0012 | 0.0058 | -5.1533 | -3.2899 | -2.7128 | -2.2839 | -1.5761 | 1.0007 | 1.0011 | 0.5758 |
| mu.p[7,3] | -1.8557 | 0.5262 | 0.0007 | 0.0018 | -2.8618 | -2.2109 | -1.8702 | -1.5089 | -0.7926 | 1.0001 | 1.0002 | 0.5758 |
| mu.p[8,3] | -1.8219 | 0.4845 | 0.0006 | 0.0018 | -2.7513 | -2.1508 | -1.8316 | -1.4983 | -0.8546 | 1.0000 | 1.0000 | 0.5758 |
| mu.p[9,3] | -2.8450 | 0.9127 | 0.0012 | 0.0055 | -5.0869 | -3.2540 | -2.6846 | -2.2537 | -1.5255 | 1.0007 | 1.0011 | 0.5758 |
| mu.p[10,3] | -2.1458 | 0.7973 | 0.0010 | 0.0025 | -3.6925 | -2.6059 | -2.1731 | -1.7218 | -0.4324 | 1.0001 | 1.0003 | 0.5758 |
| mu.p[11,3] | -2.6754 | 0.9261 | 0.0012 | 0.0050 | -4.9097 | -3.0937 | -2.5404 | -2.1037 | -1.2156 | 1.0004 | 1.0009 | 0.5758 |
| mu.p[12,3] | -2.9902 | 0.9224 | 0.0012 | 0.0061 | -5.2772 | -3.4061 | -2.8150 | -2.3765 | -1.7105 | 1.0005 | 1.0010 | 0.5758 |
| mu.p[13,3] | -2.4174 | 0.6985 | 0.0009 | 0.0026 | -3.9763 | -2.8021 | -2.3681 | -1.9773 | -1.1470 | 1.0001 | 1.0004 | 0.5758 |

| | | | | | | | | | | | |
|---|---|---|---|---|---|---|---|---|---|---|---|
| mu.p[14,3] | -2.6694 | 0.9208 | 0.0012 | 0.0050 | -4.8856 | -3.0863 | -2.5372 | -2.1003 | -1.2158 | 1.0003 | 1.0006 | 0.5758 |
| mu.p[15,3] | -2.7521 | 0.9219 | 0.0012 | 0.0054 | -5.0165 | -3.1654 | -2.6026 | -2.1690 | -1.3607 | 1.0004 | 1.0009 | 0.5758 |
| mu.p[16,3] | -2.6870 | 0.9203 | 0.0012 | 0.0048 | -4.9114 | -3.0987 | -2.5497 | -2.1157 | -1.2619 | 1.0006 | 1.0010 | 0.5758 |
| mu.p[17,3] | -2.9150 | 0.9074 | 0.0012 | 0.0057 | -5.1718 | -3.3216 | -2.7447 | -2.3166 | -1.6416 | 1.0004 | 1.0007 | 0.5758 |
| mu.p[18,3] | -2.6039 | 0.9420 | 0.0012 | 0.0045 | -4.8272 | -3.0314 | -2.4886 | -2.0447 | -1.0475 | 1.0007 | 1.0012 | 0.5758 |
| mu.p[19,3] | -2.1219 | 0.6117 | 0.0008 | 0.0017 | -3.3616 | -2.5009 | -2.1225 | -1.7384 | -0.8996 | 1.0000 | 1.0001 | 0.5758 |
| mu.p[20,3] | -2.8920 | 0.9216 | 0.0012 | 0.0059 | -5.1683 | -3.3061 | -2.7241 | -2.2892 | -1.5762 | 1.0003 | 1.0005 | 0.5758 |
| mu.p[21,3] | -2.0821 | 0.5330 | 0.0007 | 0.0015 | -3.1551 | -2.4215 | -2.0812 | -1.7363 | -1.0327 | 1.0000 | 1.0001 | 0.5758 |
| mu.p[22,3] | -2.6934 | 0.9297 | 0.0012 | 0.0052 | -4.9378 | -3.1142 | -2.5569 | -2.1182 | -1.2383 | 1.0005 | 1.0010 | 0.5758 |
| mu.p[23,3] | -2.3663 | 0.7121 | 0.0009 | 0.0024 | -3.9149 | -2.7643 | -2.3299 | -1.9299 | -1.0196 | 1.0001 | 1.0003 | 0.5758 |
| new.R3 | 1263.1464 | 28.3403 | 0.0366 | 0.0673 | 1208.3456 | 1243.9540 | 1262.8699 | 1282.1342 | 1319.4786 | 1.0000 | 1.0000 | 0.5758 |
| sigma.a | 1.4312 | 0.3496 | 0.0005 | 0.0029 | 0.8286 | 1.1903 | 1.4016 | 1.6384 | 2.2039 | 1.0006 | 1.0019 | 0.5758 |
| sigma.b1 | 0.8314 | 0.1982 | 0.0003 | 0.0010 | 0.5119 | 0.6916 | 0.8083 | 0.9452 | 1.2853 | 1.0001 | 1.0002 | 0.5758 |
| sigma.b2 | 0.3317 | 0.1003 | 0.0001 | 0.0005 | 0.1781 | 0.2596 | 0.3177 | 0.3884 | 0.5657 | 1.0000 | 1.0001 | 0.5758 |
| sigma.b3 | 0.7179 | 0.1791 | 0.0002 | 0.0009 | 0.4280 | 0.5915 | 0.6971 | 0.8213 | 1.1267 | 1.0001 | 1.0004 | 0.5758 |
| sigma.b4 | 0.3980 | 0.1116 | 0.0001 | 0.0005 | 0.2184 | 0.3185 | 0.3849 | 0.4629 | 0.6530 | 1.0000 | 1.0001 | 0.5758 |
| sigma.bp[1] | 0.9431 | 0.2689 | 0.0003 | 0.0028 | 0.5423 | 0.7502 | 0.9004 | 1.0900 | 1.5836 | 1.0012 | 1.0036 | 0.5758 |
| sigma.bp[2] | 1.2915 | 0.3914 | 0.0005 | 0.0019 | 0.6976 | 1.0150 | 1.2344 | 1.5029 | 2.2134 | 1.0002 | 1.0006 | 0.5758 |
| sigma.bp[3] | 0.7389 | 0.4181 | 0.0005 | 0.0037 | 0.2295 | 0.4411 | 0.6451 | 0.9295 | 1.7886 | 1.0012 | 1.0018 | 0.5758 |

**Table S4:** Model results for Bayesian multi-species occupancy analysis fit to 23 mammal species.

### Appendix S1

JAGS model to implement Bayesian multi-species occupancy model.

```
model{

	mu.a~dunif(-20,20)
	tau.a<-1/(sigma.a*sigma.a)
	sigma.a~dunif(0,10)

	b.leech~dunif(-20, 20)
	b.CT~dunif(-20, 20)
	b.eff~dunif(-20, 20)

	for (i in 1:3){
	mu.bp[i]~dnorm(0, 0.01)
	tau.bp[i]~dgamma(0.1, 0.1)
	sigma.bp[i]<-sqrt(1/tau.bp[i])
	}

	### fixed effects on occupancy across sites

	mu.b1~dnorm(0, 0.01)
	mu.b2~dnorm(0, 0.01)
	mu.b3~dnorm(0, 0.01)
	mu.b4~dnorm(0, 0.01)

	tau.b1~dgamma(0.1, 0.1)
	tau.b2~dgamma(0.1, 0.1)
	tau.b3~dgamma(0.1, 0.1)
	tau.b4~dgamma(0.1, 0.1)

	sigma.b1<-sqrt(1/tau.b1)
	sigma.b2<-sqrt(1/tau.b2)
	sigma.b3<-sqrt(1/tau.b3)
	sigma.b4<-sqrt(1/tau.b4)

	for (i in 1:M){
	### draw species from overall normal

	beta1[i]~dnorm(mu.b1, tau.b1)
	beta2[i]~dnorm(mu.b2, tau.b2)
	beta3[i]~dnorm(mu.b3, tau.b3)
	beta4[i]~dnorm(mu.b4, tau.b4)

	alpha[i]~dnorm(mu.a, tau.a)

	for (l in 1:3){
	mu.p[i,l]~dnorm(mu.bp[l], tau.bp[l])
	}


	### loop over all survey stations across all sites
	for (j in 1:J){
	lpsi[i,j]<-alpha[i] + beta1[i]*ELE[j] + beta2[i]*VD[j] + beta3[i]*CDL[j] + beta4[i]*LCP[j]

	psi[i,j]<-exp(lpsi[i,j])/(exp(lpsi[i,j]) + 1)
	psi.eff[i,j]<-psi[i,j]
```

```
z[i,j]~dbern(psi.eff[i,j])

for (k in 1:maxocc){
### species level detection
### survey methods: 1 = CT, 2 = brown leech, 3 = tiger leech

l.psp[i,j,k]<-mu.p[i,method[k]] + b.leech*leech_efftot[j,k] + b.CT*CT_efftot[j,k]
psp[i,j,k]<-exp(l.psp[i,j,k])/(1+exp(l.psp[i,j,k]))

p.eff[i,j,k]<-z[i,j]*psp[i,j,k] * effabs[j,k]
y[i,j,k]~dbern(p.eff[i,j,k])

### generate new data from model under consideration
new.y[i,j,k]~dbern(p.eff[i,j,k])

### calculate Freeman-Tukey residuals for real and new data to assess model fit
res[i,j,k]<-(y[i,j,k] - sqrt(p.eff[i,j,k]) )^2
new.res[i,j,k]<-(new.y[i,j,k] - sqrt(p.eff[i,j,k]) )^2
}

###sum residuals over occasions
R1[i,j]<-sum(res[i,j,])
new.R1[i,j]<-sum(new.res[i,j,])
}

###sum residuals over sites
R2[i]<-sum(R1[i,])
new.R2[i]<-sum(new.R1[i,])

#total number of occupied sites across all sites
Ntot[i]<-sum(z[i,1:J])

## species occurs or not
occt[i]<-1 - equals(Ntot[i],0)
}

R3<-sum(R2[1:23])  ### number of detected species
new.R3<-sum(new.R2[1:23])  ### number of detected species

Nst<-sum(occt[1:M])
```

### Appendix S2

To test for spatial autocorrelation we calculated Moran's I using the R package *lctools*. We calculated residuals as whether or not the species was detected at site j (detection = 1, non-detection = 0) minus the predicted probability of detecting the species at least once, given as:

$$\psi_{ij} p.tot_{ij},$$

where $\psi_{ij}$ is the occupancy probability of species *i* at site *j*, and

$$p.tot_{ij} = 1 - \prod_{k=1}^{K} 1 - p_{ijk},$$

For each species in the data set, we calculated Moran's I at a neighborhood distance of 2.5 km, following the average sampling station spacing for our study.

| **Species** | **Moran's I** <br> **(*p*)** |
|---|---|
| Annamite striped rabbit | 0.065 <br> (0.488) |
| Asian black bear | **-0.285** <br> **(0.008)*** |
| Asiatic brush tailed porcupine | -0.019 <br> (0.913) |
| Common palm civet | 0.141 <br> (0.157) |
| Crab eating mongoose | **0.214** <br> **(0.035)*** |
| Dark muntjac | 0.115 <br> (0.243) |
| Eurasian wild pig | 0.089 <br> (0.357) |
| Ferret badger | 0.059 <br> (0.526) |
| Leopard cat | 0.108 <br> (0.27) |
| Malayan porcupine | 0.114 <br> (0.246) |

| | |
|---|---|
| Marbled cat | -0.016<br>(0.934) |
| Masked palm civet | -0.056<br>(0.644) |
| Northern treeshrew | 0.125<br>(0.207) |
| Owston's civet | 0.009<br>(0.875) |
| Pangolin | -0.042<br>(0.739) |
| Pig tailed macaque | 0.155<br>(0.123) |
| Red muntjac | 0.166<br>(0.099) |
| Rhesus macaque | -0.09<br>(0.432) |
| Serow | -0.048<br>(0.697) |
| Spotted linsang | -0.057<br>(0.634) |
| Stump tailed macaque | 0.098<br>(0.316) |
| Yellow bellied weasel | 0.01<br>(0.868) |
| Yellow throated marten | -0.138<br>(0.212) |

**Appendix S3**

**Equation and application of modified Bray-Curtis dissimilarity index.**

We calculated an occupancy-based dissimilarity index for the species suite in each site by incorporating the posterior distributions of occupancy probability into the modified Bray-Curtis dissimilarity index proposed by Giacomini and Galetti (2013):

$$D(r,f) = \frac{\sum_{k=1}^{S} \omega_k \left(N_{k,r} - N_{k,f}\right)}{\sum_{k=1}^{S} \omega_k \left(N_{k,r} + N_{k,r}\right)}$$

where *D* is the index of dissimilarity of focal assemblage f with respect to a reference assemblage *r*; *S* is the total number of species in the focal (*f*) and reference (*r*) assemblages; *k* is the identification of a species; $N_{k,r}$ is the presence or absence of species k in the reference assemblage; and $N_{k,f}$ is presence or absence of species *k* in the focal assemblage. *D* ranges from 0 to 1, with 0 indicating no difference in occupancy between the focal and reference sites, and 1 indicating complete difference in occupancy.

Instead of using species presence or absence, we used the distribution of occupancy estimates from the target site as our reference assemblage, and compared the occupancy for that site with all other sites, so that $N_{k,r}$ and $N_{k,f}$ are the occupancy of species k in the reference and focal assemblage, respectively. We repeated this procedure 30,000 times to generate a distribution of *D* values and then took the mean and standard error of that distribution.
